# Designing Feature-Controlled Humanoid Antibody Discovery Libraries Using Generative Adversarial Networks

**DOI:** 10.1101/2020.04.12.024844

**Authors:** Tileli Amimeur, Jeremy M. Shaver, Randal R. Ketchem, J. Alex Taylor, Rutilio H. Clark, Josh Smith, Danielle Van Citters, Christine C. Siska, Pauline Smidt, Megan Sprague, Bruce A. Kerwin, Dean Pettit

## Abstract

We demonstrate the use of a Generative Adversarial Network (GAN), trained from a set of over 400,000 light and heavy chain human antibody sequences, to learn the rules of human antibody formation. The resulting model surpasses common *in silico* techniques by capturing residue diversity throughout the variable region, and is capable of generating extremely large, diverse libraries of novel antibodies that mimic somatically hypermutated human repertoire response. This method permits us to rationally design *de novo* humanoid antibody libraries with explicit control over various properties of our discovery library. Through transfer learning, we are able to bias the GAN to generate molecules with key properties of interest such as improved stability and developability, lower predicted MHC Class II binding, and specific complementarity-determining region (CDR) characteristics. These approaches also provide a mechanism to better study the complex relationships between antibody sequence and molecular behavior, both *in vitro* and *in vivo*. We validate our method by successfully expressing a proof-of-concept library of nearly 100,000 GAN-generated antibodies via phage display. We present the sequences and homology-model structures of example generated antibodies expressed in stable CHO pools and evaluated across multiple biophysical properties. The creation of discovery libraries using our *in silico* approach allows for the control of pharmaceutical properties such that these therapeutic antibodies can provide a more rapid and cost-effective response to biological threats.

## INTRODUCTION

Antibodies are an important class of biologics-based therapeutics with clear advantages of specificity and efficacy.^1,2,3^ The high cost and long development times, however, present key challenges in the accessibility of monoclonal antibody therapeutics.^4,5^ To quickly respond to known and new pathogens and disease, and to provide affordable, high quality treatment to patients around the globe, a molecule must be designed for activity; but it must also be made developable and safe for patients. Adding to their overall cost and process time, many antibodies suffer from poor yields or require individually customized processing protocols or formulations because their biophysical properties cause them to aggregate, unfold, precipitate, or undergo other physical modification during processing into a drug product.^6–13^ Even with the significant amounts of research being put into discovering pharmacologically-active antibodies and understanding their physical and biological behavior, they remain challenging to identify for given diseases or pathogens and to optimize for developability.

Discovery of therapeutic antibodies frequently involves either display methodologies or B-cells isolated from humans or animals that have been exposed to an antigen or a disease target of interest.^14,15^ Although B-cell isolation and deep sequencing workflows have improved over the years in regards to cost, labor, and speed, there are still inherent limitations when considering the process as an antibody discovery platform. A sufficient immunological response is required from the specific subjects being used, and due to the low number and diversity of subjects, there can be insufficient antibody sequence diversity that is expressed. There is also the challenge of overcoming B-cell-driven survival against specific epitopes in which therapeutically viable epitopes are not utilized by an immune response when they are out-competed by a dominant binding epitope, leading to an antibody panel focused on a limited epitope. The library approach can provide a search across a wider range of sequence space, but most examples of synthetic libraries result in a sequence profile which is quite different from those expressed by the human immune system. In both cases, there is little to no ability to control the chemical, biophysical, or biological characteristics of the identified candidates. As a result, discovered antibodies frequently have the aforementioned features seriously complicating their developability and stability.

A recent synthetic library approach implements random mutagenesis in which specific residues are allowed to vary in type following statistical rules for frequency of appearance by location in the antibody (commonly known as positional frequency analysis, PFA).^16,17,18^ PFA and other related methods^19,20,21^ do not take into account any interactions between residues except to the extent that such interactions limit the expressibility of the protein. While this widely explores the sequence space, it ignores how residue types interact to form stabilizing features such as hydrogen or ionic bonds. Random assignment is also done without consideration of the characteristics of the final antibody entity, leading to some that have unusual and potentially problematic protein surface features.

Another drawback to most synthetic library approaches is that they focus solely on the complementary-determining regions (CDRs) of the antibodies. While the CDRs are the most critical portion of the antibody variable region in determining binding interactions, many Kabat-defined CDR positions are part of the core immunoglobulin (Ig) fold, and many of the framework residues can also play an important role in direct antigen binding, stability of the molecule, and CDR orientation.^6^ By limiting mutations to the CDRs, existing libraries neglect the possibility of improved bioactivity and developability afforded by some framework mutations.

Even with an identified therapeutic antibody, improving that antibody’s production and purification behavior through sequence modification can be challenging. While many papers have been published trying to develop a predictable connection between an antibody’s sequence and/or computed molecular structure and the molecule’s various physical characteristics, the connection is elusive as it involves complex nonlinear interactions between the constituent amino acid residues.^11,22–37^ Frequently, such work involves an exceptionally small number of molecules, frequently under 200 and often under 50, from a non-diverse set of sequences - a small number of parental sequences, several parents with a small number of highly-related sequence variants, or a single antibody with mutational scanning. Such approaches give information on an individual antibody or small group, but are highly unlikely to generalize the complexity of residue interactions to other antibodies. Such understanding requires exploration of the wider hyperdimensional space of antibody sequences. Computational approaches used to optimize molecular behavior also frequently ignore whether the revised molecule remains similar to human antibodies. That assessment is left to expensive *in vitro* studies.

Deep learning offers one route to better capture the complex relationships between sequence and protein behavior and has been the focus of many recent publications.^38–42^ Within the context of discovery and libraries, the generative models such as Generative Adversarial Networks (GANs)^43,44^ and autoencoder networks (AEs)^45^ are of particular interest as they have been shown to be viable for generating unique sequences of proteins^46,47^ and nanobodies^48^ and antibody CDRs^49^. But these efforts focus on short sequences of proteins or portions of antibodies. Use of these approaches in the full antibody sequence space entails a unique set of challenges for machine learning models.

Antibodies derive from different germline backgrounds, are much larger in size, and are composed of multiple chains, leading to a more complex sequence and structural space. More complexity in a machine learning setting generally requires more data to resolve. However, sequence data, with associated experimental data, is more limited for antibodies and is far more costly to come by than small molecules.

Here, we present the Antibody-GAN, a new synthetic approach to designing a novel class of antibody therapeutics which we term “humanoid” antibodies. The Antibody-GAN uses modified Wasserstein-GANs^50^ for both single-chain (light or heavy chain) and paired-chain (light and heavy chain) antibody sequence generation. These GANs allow us to encode key properties of interest into our libraries for a feature-biased discovery platform. Our Antibody-GAN architecture captures the complexity of the variable region of the standard human antibody sequence space, (2) provides a basis for generating novel antibodies that span a larger sequence diversity than is explored by standard *in silico* generative approaches, and (3) provides, through transfer learning (continued training of a model with a subset of data with specific desirable characteristics), an inherent method to bias the physical properties of the generated antibodies toward improved developability and chemical and biophysical properties.

We demonstrate the GAN library biasing on such properties as a reduction of negative surface area patches, identified as a potential source of aggregation, thermal instability, and possible half-life reductions,^51^ and away from MHC class II binding, which may reduce the immunogenicity of the generated antibodies.^52–56^ We show, additionally, library biasing to a higher isoelectric point (pI) to reduce aggregation and prevent precipitation in therapeutic formulations, and towards longer CDR3 lengths which can increase diversity and has been known to create more effective therapeutics for a class of targets.^57^

To demonstrate the viability of the Antibody-GAN to generate humanoid antibody sequences, the GAN was used to generate a proof-of-concept validation library of 100k sequences from 4 germline subgroups. These sequences were generated using two single-chain GANs (each trained on a set of 400,000 heavy or light chain sequences from human-repertoire antibodies).^58^ The GAN sequences were expressed as antibody antigen binding fragments (Fabs) in phage. Two of the less-represented germline subgroups were optimized for germline agreement using transfer learning. From this initial library, we present the sequences, structure, and biophysical properties of two antibodies with divergent surface patch features which were expressed in stable Chinese hamster ovary (CHO) cells.

## RESULTS

### Generative Adversarial Networks for Antibody Design

The general Antibody-GAN architecture is depicted in **Figure 1A** in which a set of real training variable-region (Fv) antibody sequences are fed to a discriminator of the GAN along with the output of the generator. The generator takes a vector of random seeds as input, and outputs a random synthetic antibody sequence. During training, the discriminator is progressively trained to attempt to accurately distinguish between the real and the synthetic sequences and the generator is progressively trained to produce synthetic sequences that cannot be distinguished from the real human repertoire sequences in the training set. After initial training of the Antibody-GAN from a general training set, transfer learning can be used to bias the GAN towards generating molecules with specific properties of interest. The details of the underlying neural network architecture are given in the supplementary information (**Figure S1**).

**Figure 1.**
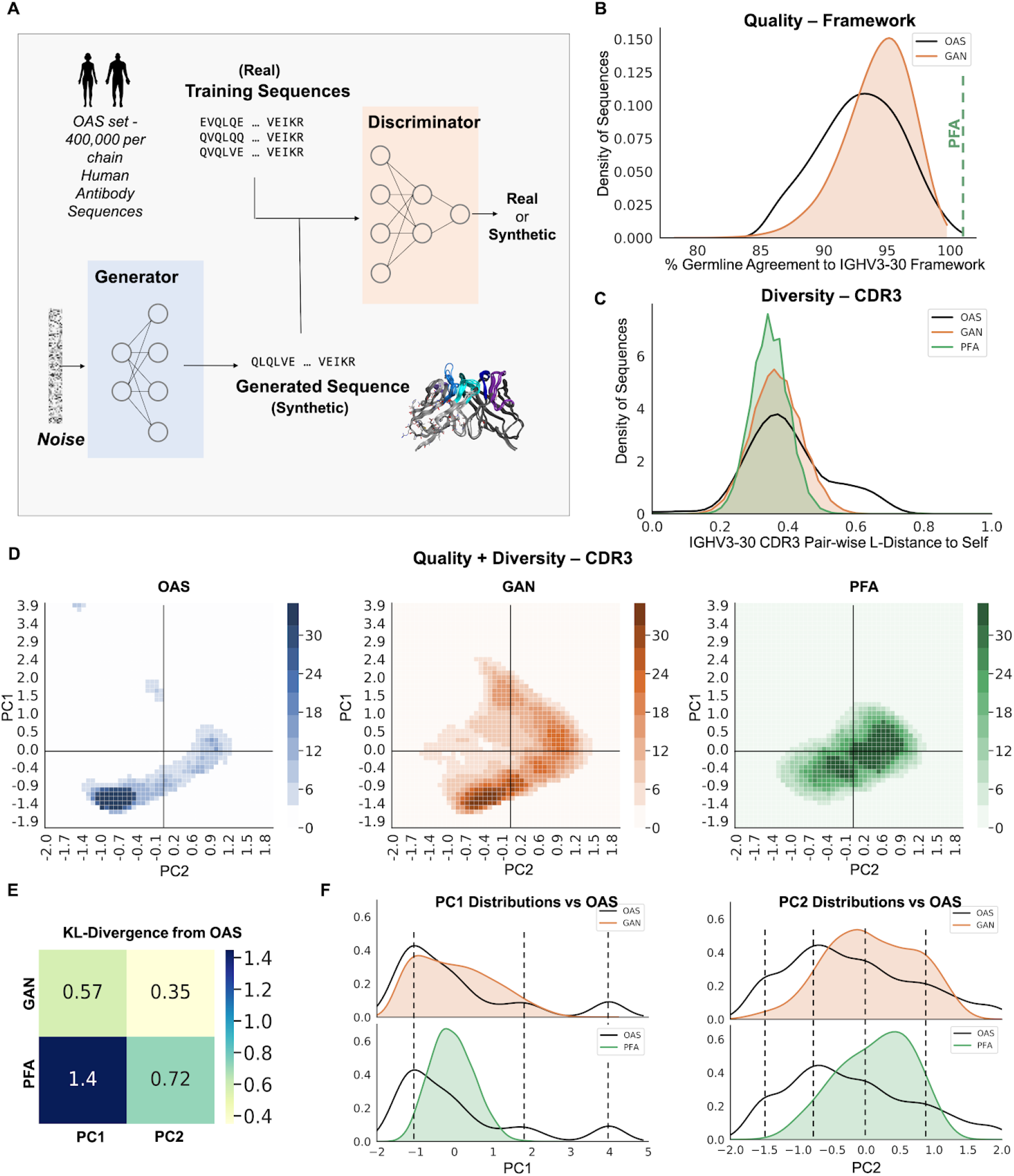
GAN-generated human antibody sequences. **A)** Schematic representation of the Antibody-GAN architecture; **B)** Representative percent germline agreement for framework residues from the training (OAS) and unbiased GAN (GAN) for 10,000 generated sequences per set. All PFA sequences have 100% germline agreement (green dashed-line); **C)** Representative diversity in heavy chain CDR3, as Levenshtein distance calculated within sets of 10,000 sequences from each of: the OAS training set, GAN-generated molecules, and PFA-generated molecules; **D)** Quality and diversity of heavy chain CDR3 as captured in the scores of a principal component analysis (PCA). The GAN set shows more similar density and coverage in the PCA space to human repertoire (OAS) than PFA; **E)** the KL-divergence of each principal component for GAN and PFA, compared to OAS, shows lower divergence of GAN from OAS than PFA; **F)** The independent distributions of PC1 and PC2 of GAN and PFA compared to OAS again show GAN reproducing human repertoire better than PFA.

As a demonstration of the general architecture and training approach, an Antibody-GAN was trained using a set of 400,000 human-repertoire sequences, per chain, randomly selected from the Observed Antibody Space project (OAS).^58^ Prior to training, the sequences were all structurally aligned using the AHo numbering system which enables direct comparison of residues at the same structural position across the dataset.^59^ This greatly simplifies the relationships that the GAN must capture both for generation and discrimination. Additional training details are provided in the methods section.

In **Figure 1B**, sequences from the Antibody-GAN (GAN), the OAS training set, and a set of sequences with 100% germline framework and PFA-generated CDRs (PFA) were compared by selecting a random set of 10,000 sequences, all classified as germline HV3-30, from the training set and all three synthetic sets. These were evaluated on the distribution of percent germline agreement of the framework residues. Within the human repertoire, deviations from framework germline agreement arise from the sequences having undergone somatic hypermutation during B-cell maturation. Molecules from the Antibody-GAN model (GAN) deviate from germline much like the OAS. Note that the PFA set uses an exact germline framework; as such, the germline agreement is always 100%.

The diversity of the heavy variable (HV) CDR3 was used as an indicator of the diversity of binding paratopes within a given set and was assessed using (1) pairwise Levenshtein distances calculated from only the HV CDR3 residues in all three sets (**Figure 1C**), and (2) the scores from the first two components of a principal component analysis (PCA) model on the aligned HV CDR3 sequences from the OAS, GAN, and PFA data (**Figure 1D**) (See supplementary **Figures S2** and **S3** for kappa variable (KV) germline agreement, CDR3 Levenshtein distance, and PCA plots). The distribution of pairwise Levenshtein distances between HV CDR3 sequences across the three sets of antibodies are shown in **Figure 1C**, where a distance of 0 indicates identical sequences and larger values indicate more dissimilar sequences. The OAS set in general shows the greatest diversity in HV, however, the GAN and PFA sets have similar diversity to the main peak in OAS, with the GAN exhibiting slightly larger diversity than PFA.

The PCA scores biplots in **Figure 1D** show that the distribution of sequence variability in the OAS human repertoire set (left plot) is more similar to those seen in the GAN set (middle plot), than the distribution in the PFA set (right plot), which diverges significantly from the other two sets, particularly in the high-density regions of the plots. The explained variance of PCA components 1 and 2 are 10% and 4%, respectively. While these are only small portions of the overall variance in the HV CDR3, they do represent the largest covarying relationships between the HV CDR3 residues and indicate that the Antibody-GAN approach captures significant relationships in human repertoire HV CDR3 that the PFA approach does not. The KL-divergence is a measure of how different two distributions are from each other, with a value of 0 indicating identical distributions and values tending away from 0 indicating more divergent distributions. The KL-divergence of the distribution over PC1, the component that captures most of the variance in CDR3, for the OAS and GAN sets is 0.57 (**Figure 1E**). The KL-divergence of PC1 for the OAS and PFA sets is 1.45. **Figure 1F** shows these distributions for PC1 and PC2 for both the GAN and the PFA sets, relative to OAS. The PFA set shows notably more divergence from the OAS and GAN sets and raises the question of how well the PFA approach reproduces human paratopes, as well as the diversity of these paratopes.

### Bias and Control of Antibody Discovery Libraries

Our generative, deep-learning approach to humanoid antibody library generation not only results in antibody libraries that are more human-like than existing synthetic library approaches, but it also allows us to control the features of our libraries. **Figure 2** shows the distributions of several properties for different libraries as they compare to OAS, with each distribution containing a representative sample of 10,000 sequences. A subset of these libraries were generated using a deep-learning technique known as transfer learning, which biases networks from the general Antibody-GAN-learned properties towards specific features of interest, such as those shown in **Figure 2**. The corresponding heatmap shows the difference, in percent of sequences in a given bin, between each library and OAS.

**Figure 2.**
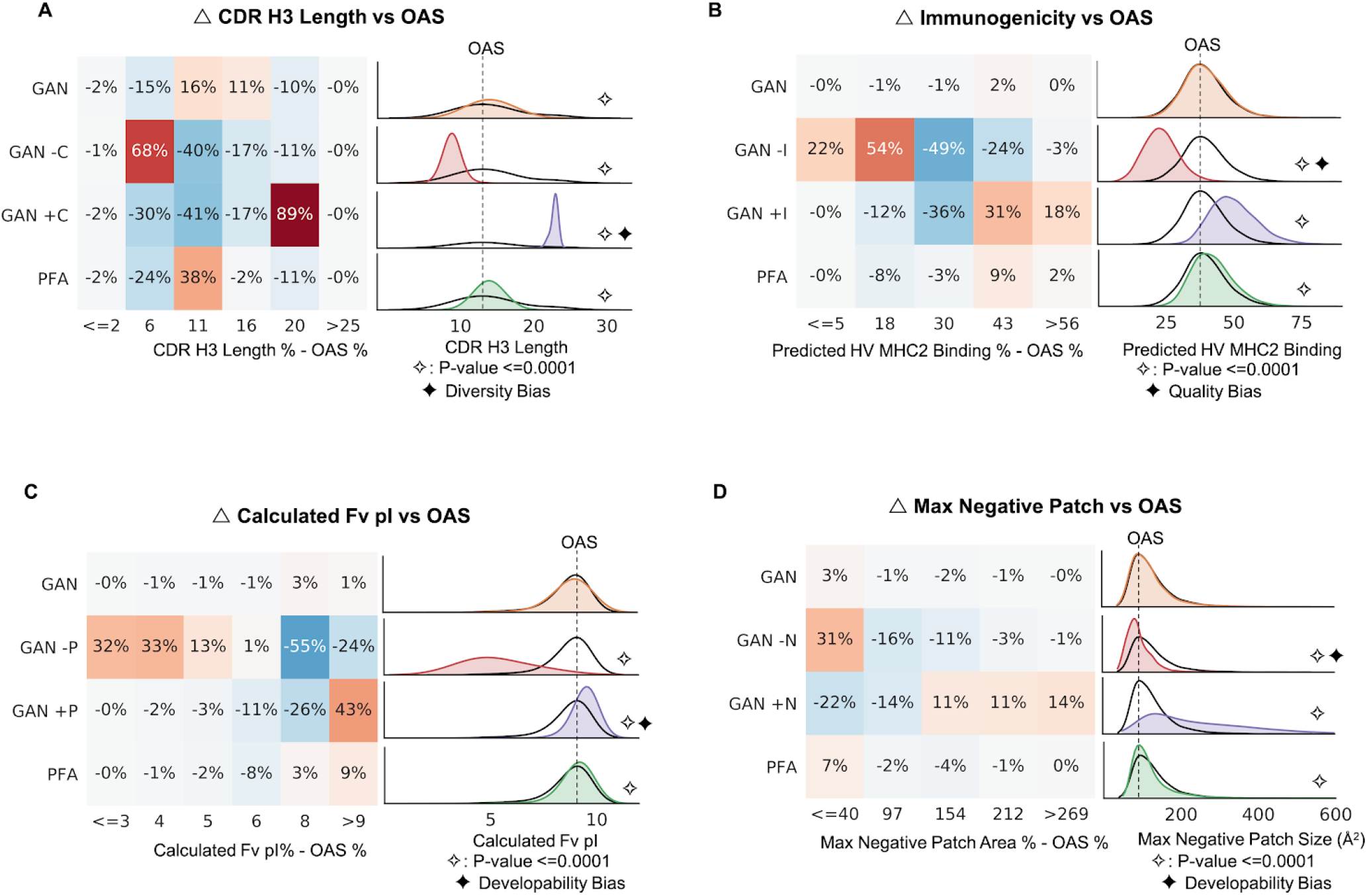
Bias and control of Antibody-GAN library features using transfer learning. **A)** Representative heavy-chain CDR3 length, as compared to OAS, from different sets of 10,000 sequences each: the unbiased Antibody-GAN (GAN), Antibody-GANs transfer learned to small (GAN −C) and to large (GAN +C) CDR H3 lengths, and PFA-generated (PFA) molecules; **B)** Representative predicted heavy chain MHCII binding, as compared to OAS, for the unbiased Antibody-GAN (GAN), Antibody-GANs transfer learned to lower (GAN −I) and to higher (GAN +I) predicted MHCII binding, and PFA-generated (PFA) molecules; **C)** Representative calculated isoelectric point (pI) on the Fv across both heavy and light chains, as compared to OAS, for the unbiased Antibody-GAN (GAN), Antibody-GANs transfer learned to low (GAN −P) and to high (GAN +P) pI, and PFA-generated (PFA) molecules; **D)** Representative maximum negative patch size across both heavy and light chains, as compared to OAS, for the unbiased Antibody-GAN (GAN), Antibody-GANs transfer learned to small (GAN −N) and to large (GAN +N) max negative surface patch size, and PFA-generated (PFA) molecules.

**Figure 2A** demonstrates biasing on the length of the primary binding paratope, CDR H3, of an antibody sequence. We compare 4 libraries to OAS: the baseline Antibody-GAN (GAN) from **Figure 1**, Antibody-GANs transfer learned to small (GAN −C) and to large (GAN +C) CDR H3 lengths, and the PFA-generated library (PFA) from **Figure 1**. The baseline Antibody-GAN library shows a total 27% difference from OAS in its CDR H3 distribution. Though still significantly different, it more closely reproduces the OAS distribution over CDR H3 than PFA (38% difference) or the other two intentionally biased libraries. The GAN −C library was generated by a model transfer-learned on a small subset of about 1,000 sequences from the GAN library which had CDR H3 lengths of less than 12 and resulted in a library with 68% shift to shorter CDR H3 sequences. The GAN +C was similarly transfer-learned on approximately 1,000 sequences from the GAN library which had CDR H3 lengths of >22, creating a very significant 89% bias towards longer CDR H3 sequences (**Figure S9**). By creating antibodies with longer CDRs, and therefore more residues to vary, the GAN +C library also inherently biases towards diversity. Antibodies with long CDR H3s have also been shown to have better success as therapeutics for diseases such as human immunodeficiency virus (HIV),^57^ and may be useful as a discovery sub-library for targets that may require such long, exposed paratopes.

**Figure 2B** shows biasing on immunogenicity of the heavy chain (see **Figure S4** for light chain biasing) using a major histocompatibility class II (MHCII) binding score derived from *in silico* peptide fragment-MHCII binding affinity predictions. We use an in-house machine learning predictor for peptide-MHCII binding (**Figure S5**) similar in effect to the binding prediction tools provided by the Immune Epitope DataBase (IEDB).^52,54,55,56,60^ Peptide-MHCII binding is the first step in the T-cell-mediated immune response and the clearest handle available for practically mitigating immunogenicity risk.^7,22^ The GAN library, with only a 2% difference from OAS in predicted immunogenicity, is statistically indistinguishable (at p<0.0001) from the human repertoire training set, whereas PFA shows a statistically significant 11% shift towards higher immunogenicity. The GAN −I library, using a similar transfer learning approach to the one described above, shows a total 76% shift to lower predicted MHCII binding than human repertoire. Reduced MHCII binding is presumed to reduce the likelihood of immunogenic response as the binding is a necessary first step in that response. The resulting biased GAN −I should generate molecules with lower chance of immunogenic response. This large bias to sequences of lower immunogenicity is a significant bias towards higher quality antibody therapeutics, and could result in a library of safer treatments for patients.

The extent to which this biasing corresponds to lower immunogenicity will largely depend on the quality of the model used to choose the transfer samples. As a control condition, the GAN +I library shows 49% bias towards increased MHCII binding. While such a higher-immunogenicity biased model would not usually be of interest in developing a library, it could provide a means to generate molecules to help validate the underlying MHCII binding model, yet again highlighting the utility of a GAN method as a tool to explore molecular and therapeutic space.

For antibody therapeutics, the isoelectric point (pI, the pH at which the molecule is neutral) is a key measure of developability since a pI near the formulation pH may lead to high viscosity and aggregation or precipitation.^61–64^ A slightly acidic pH generally results in a higher overall charge leading to more recent formulations centered around pH 5.5.^65^ To remain stable in solution, therapeutic antibodies would ideally need to have a pI of greater than 8.5 for the overall molecule. The GAN library provides a distribution of pI for the Fv portion of the antibody that is statistically indistinguishable from OAS (**Figure 2C**), and the PFA library creates a small 11% bias towards higher Fv pI. We show, with the GAN −P library, that we can bias the library with a 79% shift to lower Fv pI via transfer learning. The GAN +P library, however, shows a 43% increase in sequences with a calculated Fv pI greater than 9, resulting in likely a significant bias towards developability.

Large surface patches in antibody therapeutics have been linked to developability issues such as aggregation, thermal instability, elevated viscosity, and increased clearance rate,^23,33,37^ but also to improvement of specificity,^31^ particularly when the patches are related to charge. As such, biasing a library towards larger or smaller patches might have beneficial effects. They also serve as an example of generic biasing models towards desired structural properties. **Figure 2D** shows biasing on the maximum negative surface patch area of a molecule, calculated using structure-based homology modeling. Large negative patches have been shown to increase antibody viscosity at high therapeutic concentrations.^23,26^ Once again, the GAN library is statistically equivalent to OAS in maximum negative patch size with only a 3% difference, demonstrating the model’s ability to capture human repertoire. The PFA library maintains a small but significant 7% shift to lower negative surface patch area. The GAN −N library shows that we can intentionally shift our library towards smaller negative surface patches and away from known developability issues with a 31% bias, as shown in GAN −N. The GAN +N library shows that we can also shift in the other direction with a 36% bias towards larger negative patches. Structure-based properties like surface patch can be more difficult to bias than sequence-based ones due to (1) the non-Gaussian distribution of the property and (2) the added layer of abstraction and complexity away from sequence. These issues can likely be resolved by increasing the number of sequences in the transfer learning training set by, for example, iteratively training and sampling. For more complex properties layers can be added to the model itself during transfer-learning.

### Combinatorial Library Design and Expression of Diverse Germlines

The synthesis of a diverse, *de novo* antibody discovery library comprising specific sequences can be costly. Such specific sequence targeting cannot be done with standard codon degeneracy approaches. To greatly reduce this cost, we used a chain-oriented approach to our library design, combinatorially combining heavy and light chains that are created with specific amino-acid sequences designed by the Antibody-GAN rather than designing each Fv individually. The Antibody-GAN architecture (**Figure S1**) is designed to be modular. After training with paired-chain Fv sequences, the heavy chain generator and light chain generator can be separately used to generate single-chain sequences, such that any independently generated heavy chain should pair with any independently generated light chain to create a full Fv which maintains the library’s intended features. All Antibody-GAN and transfer-learned Antibody-GAN libraries in **Figure 2** were generated from models trained in this manner.

It is also possible to split apart the Antibody-GAN model, initially, into single chain models. These must be trained on single chain sequences and may be useful when creating diverse libraries of varying germlines, when there is no property of interest associated with the full Fv for which we want to bias. Because there are few public data sets providing developability, expression, stability, and other properties on paired-chain sequences, we choose to synthesize a naive, unbiased initial discovery library to express in phage. Our goal for this first library is to reproduce human repertoire. In doing this, we will also create a data set which can greatly inform biasing of future libraries. As such, the subsequent GAN libraries and molecules were generated using the single chain version of the Antibody-GAN.

**Figure 3A** shows the distribution of the 15 most represented heavy chain germlines in the OAS training set. **Figure 3B** shows the same for kappa light chain. For our initial library, we selected heavy chain germlines IGHV3-30 and IGHV1-2 to pair combinatorially with light chain germlines IGKV3-20 and IGKV1-39. The number of training set examples for IGHV1-2 and IGKV1-39 is lower than for the other two germlines, such that there are not enough examples to train a model of sufficient quality. The problem is compounded for other germlines with even fewer training examples. This can be remedied again by using transfer learning.

**Figure 3.**
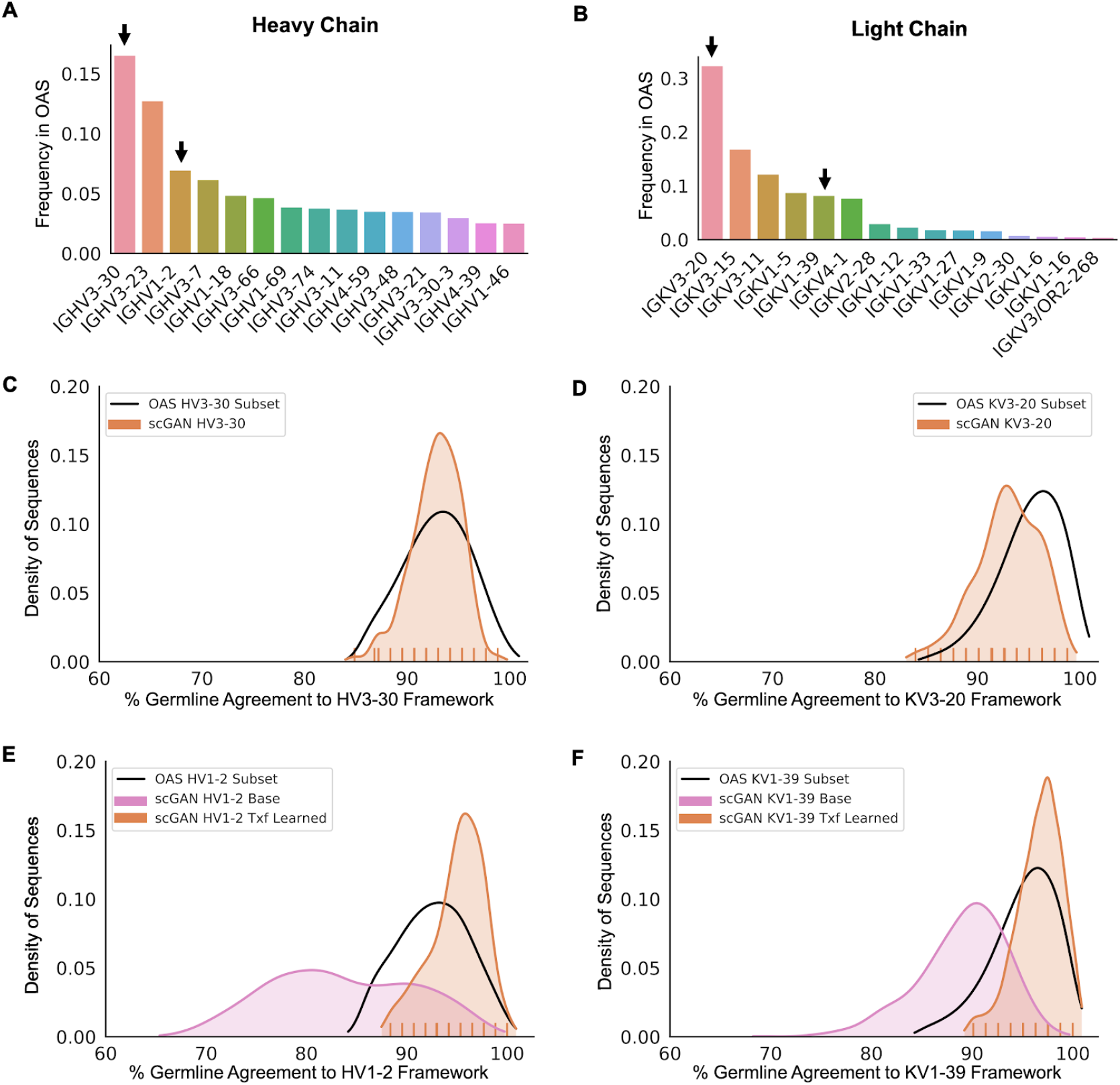
Transfer learning on less-represented germlines for a diverse library. **A)** Distribution of the 15 most represented heavy chain germlines in the OAS training set, with arrows indicating germlines used for the test library; **B)** Distribution of the 15 most represented kappa chain germlines in the OAS training set; **C)** Representative percent germline agreement for HV3-30 framework residues from the OAS and the single chain HV3-30 GAN; **D)** Representative percent germline agreement for KV3-20 framework residues from the OAS and the single chain KV3-20 GAN; **E)** Representative percent germline agreement for HV1-2 framework residues from the OAS and the single chain HV1-2 GAN, before (pink) and after (orange) transfer-learning; **F)** Representative percent germline agreement for KV1-39 framework residues from the OAS and the single chain KV1-39 GAN, before (pink) and after (orange) transfer learning.

**Figures 3C** and **3D** show the representative distributions of percent germline agreement in framework to HV3-30 and KV3-20, respectively. Because these two germlines are well-represented in the OAS training set, the models generate sequences of sufficient framework quality. **Figure 3E** and **3F**, however, show more divergence from OAS in framework quality for the less-represented HV1-2 and KV1-39 germlines, respectively, when generated by a base model without transfer learning. Only when the model is transfer-learned, allowed to continue training on only the germline subgroup of interest, is it then able to generate sequences with framework quality more closely matching OAS for HV1-2 and KV1-39. **Figure 3** shows the final synthetic GAN sub-libraries for each single-chain germline.

While a full-scale production library might contain 10,000 or more individual single-chain sequences from each germline combined combinatorially to form billions of molecules, a proof-of-concept miniature library was created by selecting 158 sequences from each of these four germlines and combining them combinatorially to assemble a library of around 100,000 total sequences. The tick marks in each distribution in **Figure 3** show the germline agreement of the sequences selected for this example library.

**Figure 4** shows Fab fragment display levels for our 4 germline-paired sub-libraries, each containing ~25,000 GAN-generated sequences. Display levels were estimated by capturing serial dilutions of purified phage on ELISA plates coated with anti-human Fab and detecting with anti-M13 antibodies conjugated to HRP. **Figure 4A** shows the average display levels, normalized for total phage concentration, of IgG Fab from each of the sub-libraries in polyclonal phage. A slight bias for higher expression can be seen at higher concentrations for those germline sub-libraries which contain the KV1-39 sub-library. Whether or not this difference is actually significant and related to higher tolerability of KV1-39 sequences, or represents differential binding of the anti-human Fab capture antibody used in the ELISA, is an area of future studies.

**Figure 4.**
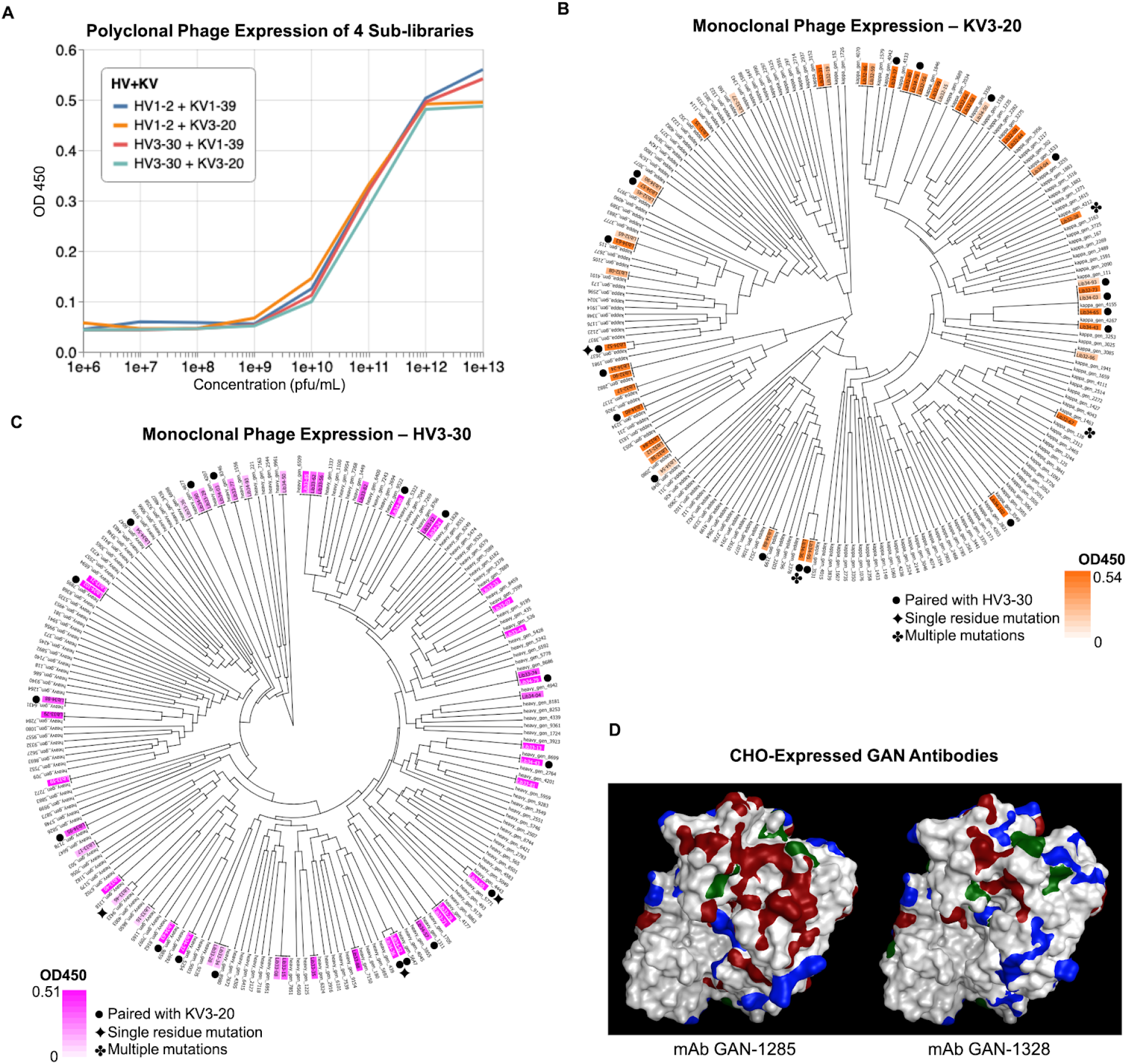
Combinatorially-assembled GAN sequences expressed in phage and CHO. **A)** ELISA showing Fab expression of 4 GAN sub-libraries from polyclonal phage. **B)** Clading of variable region protein sequences of select phage colonies expressing the KV3-20 library as determined by DNA sequencing. Orange highlighting indicates degree of expression level of selected Fabs as determined by ELISA. Sequences that were paired with HV3-30 heavy chains(●) are as marked, otherwise they were paired with HV1-2 heavy chains. Sequences with single amino acid substitutions(✦) or multiple amino acid substitutions(✤) as compared to their intended GAN design are as marked. **C)** Clading of variable region protein sequences of select phage colonies expressing the HV3-30 library as determined by DNA sequencing. Magenta highlighting indicates degree of expression level of selected Fabs as determined by ELISA. Sequences that were paired with KV3-20 light chains(●) are as marked, otherwise they were paired with KV1-39 light chains. Sequences with single amino acid substitutions(✦) or multiple amino acid substitutions(✤) as compared to their intended GAN design are as marked. **D)** 3D homology structure model of two GAN molecules expressed in CHO. Negative surface patches are shown in red, positive patches in blue, and hydrophobic patches in green.

To confirm the expressed Fab are indeed the designed, *de novo* sequences, we selected and sequenced ~30 colonies expressed in monoclonal phage from each of the 4 sub-libraries. **Figure 4B** presents a variable-region clading of the selected sequences from the two sub-libraries expressing KV3-20 light chain to the 158 GAN-designed KV3-20 sequences. This shows primarily 1) our selection of colonies was random and provides good coverage of the space of designed sequences and 2) only a small fraction of the expressed sequences contained any amino acid mutations relative to the GAN-designed sequences; most matched our synthetic designs exactly. **Figure 4B** also shows the optical density (OD450) for each of these phage-expressed sequences and whether the particular sequence was paired with a heavy chain from the HV3-30 library. Phage library sequences that are not marked as being paired with a heavy chain from the HV3-30 library were paired with a heavy chain sequence from the HV1-2 library.

**Figure 4C** presents a similar clading of the selected sequences from the two sub-libraries expressing HV3-30 heavy chain. The same observations can be made with this set. The selected sequences span the design space well, and show even fewer amino acid mutations and more exact matches to the *de novo* GAN-designed sequences than the KV3-20 set. Again, expression is shown for the sampled colonies and it is noted whether those sequences were paired with the KV3-20 light chain. Phage library sequences that are not marked as being paired with a light chain from the KV3-20 library were paired with a light chain sequence from the KV1-39 library. (See **Figures S6** and **S7** for similar clading on HV1-2 and KV1-39 sets).

A further subset of antibodies were selected from the HV3-30/KV3-20 sub-library to be expressed in stable CHO pools for biophysical analysis. **Figure 4D** presents the 3D structure of two such GAN antibodies selected, in particular, for their interesting patch properties. The molecule mAb GAN-1285 was selected for its very large negative surface patch of ~600 Å^2^, shown in red. Molecules with such a large maximum negative patch are relatively uncommon in the base Antibody-GAN distribution (**Figure 2D**), but are interesting to investigate for developability purposes. The molecule mAb GAN-1328, by contrast, has a maximum negative surface patch of ~ 140 Å^2^. The positive and hydrophobic surface patches for both molecules are marked in blue and green, respectively. **Figure 5** presents the aligned sequences of these two selected GAN-derived antibodies, showing framework mutations from germline.

**Figure 5.**
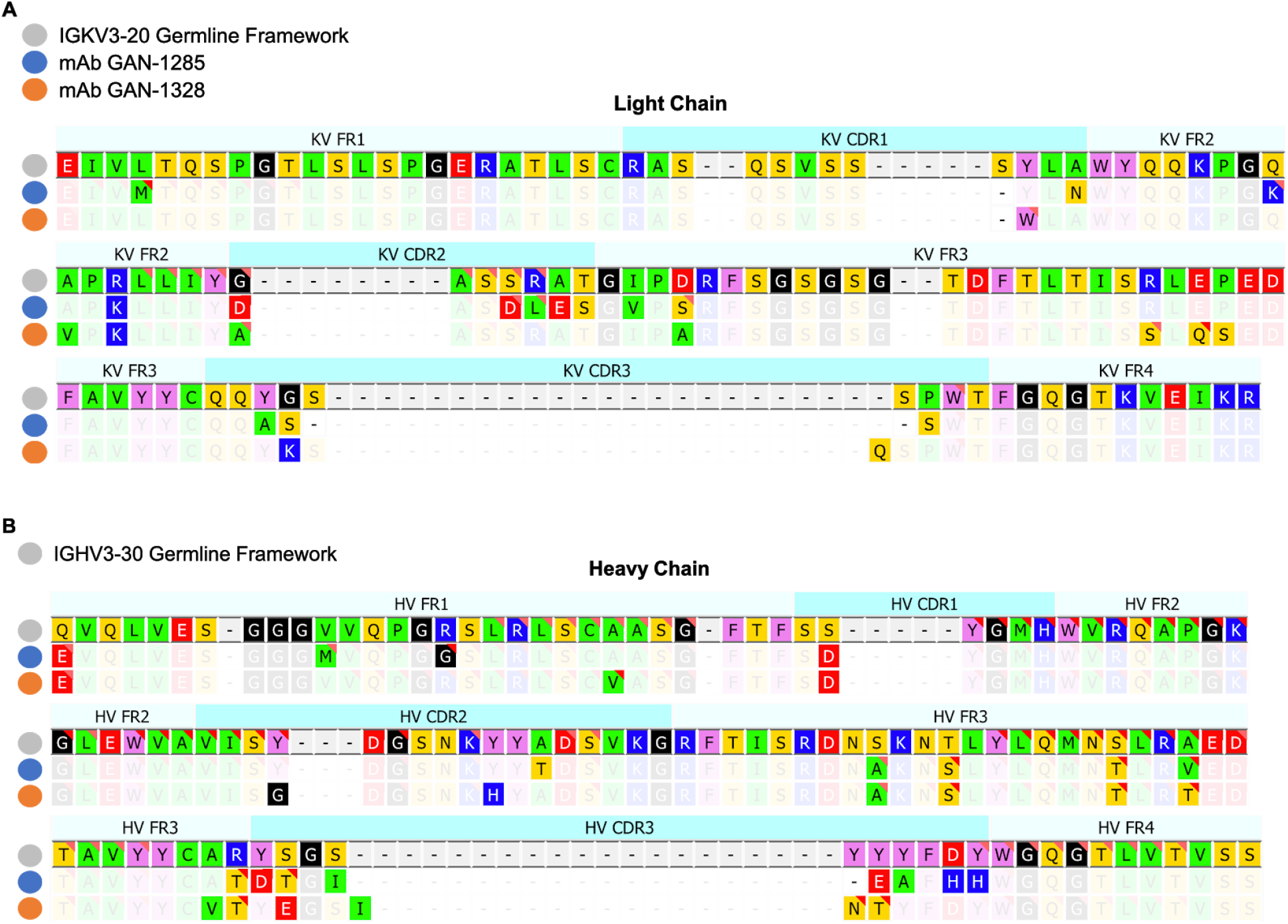
Sequence alignment of select GAN antibodies. **A)** Light chain sequence alignment to IGKV3-20 framework of two GAN molecules expressed in CHO, with faded consensus sequence; **B)** Heavy chain sequence alignment to IGHV3-30 framework of two GAN molecules expressed in CHO, with faded consensus sequence. Deviation from the germline framework, mimicking somatic hypermutation, can be seen as well as differences in CDR regions for both the light chain and heavy chain sequences in both molecules.

### Biophysical Validation of CHO-Expressed GAN Antibodies

For the purpose of validation of our GAN approach and to interrogate the interesting property of maximum negative surface patch, we present the biophysical data of mAb GAN-1285 and mAb GAN-1328 (**Figures 4D, 5**) after stable CHO expression and purification. **Figure 6** shows the behavior of these two antibodies across four key assays in our platform: differential scanning fluorimetry (DSF), self-interaction nanoparticle spectroscopy (SINS), polyethylene glycol (PEG) solubility, and size-exclusion chromatography (SEC). These assays are commonly used to assess the stability and developability of therapeutic antibodies.^66,67^

**Figure 6.**
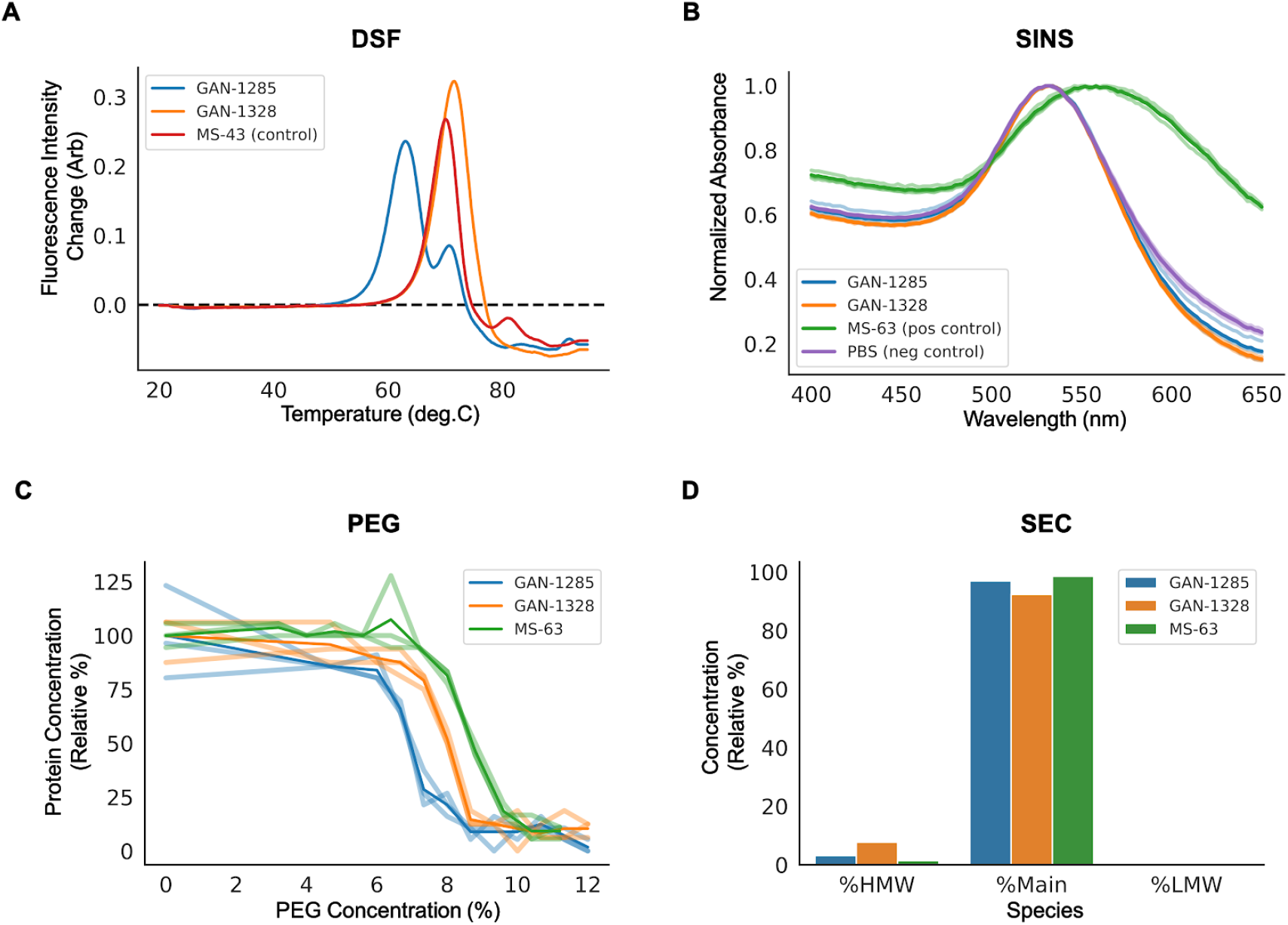
Biophysical properties of select GAN antibodies. **A)** Differential scanning fluorimetry (DSF) results for 3 CHO-expressed antibodies, two GAN-generated molecules mAb GAN-1285 (large negative patch) and mAb GAN-1328 (small negative patch), and a platform control MS-43; **B)** Self-interaction nanoparticle spectroscopy (SINS) for the two GAN antibodies, a positive platform control, MS-63, and a negative phosphate-buffered saline (PBS) control, each with three replicates and mean results as bolded lines; **C)** Polyethylene glycol (PEG) solubility results, with replicates (mean results represented by bolded lines), for the two GAN antibodies and the positive platform control **D)** Size-exclusion chromatography (SEC) results of high molecular weight (HMW), main antibody, and low molecular weight (LMW) species for the two GAN antibodies and a platform control.

DSF assesses the temperature at which certain regions of the antibody begin to unfold. More stable, and thus more developable antibodies tend to have one or more regions unfold at higher temperatures and have a higher first-unfolding transition temperature. **Figure 6A** shows the DSF results for mAb GAN-1285 and mAb GAN-1328, as well as a platform control antibody, MS-43. The identical constant regions of these three molecules all show an unfolding event at around 72 °C, presumed to be the IgG CH2 region. The molecule with a very large negative surface patch, mAb GAN-1285, shows much lower thermal stability, with an initial unfolding event near 60 °C. This is consistent with the notion that negative surface patches are known to be related to thermal instability.

The SINS assay (**Figure 6B**) is commonly used to interrogate whether a molecule will self interact, leading to issues in the manufacturing process as well as possible viscosity and filterability issues. Interestingly, the two GAN molecules exhibited the same SINS profile as the PBS negative control, indicating a low propensity to self interact, particularly compared to the positive control molecule, MS-63, known to have high self-interaction behavior.

The remaining assays, PEG solubility (**Figure 6C**) and SEC (**Figure 6D**), show that both antibodies are reasonably soluble, and display relatively low amounts of high molecular weight (HMW) formation, although there are potentially significant differences between the two antibodies in both assays.

## DISCUSSION

We describe here a new class of *de novo* human antibodies derived *in silico*, which we refer to as “humanoid”, owing to their explicit requirement that generated sequences must mimic human antibody sequence patterns. While antibodies have excellent antigen specificity and have often been adapted as scaffolds for therapeutic applications, B-cells do not undergo selective pressure *in vivo* to produce antibodies which have ideal biotherapeutic characteristics. To reduce the cost of development and greatly increase the time-to-response for known or new diseases and pathogens, a discovery library must contain therapeutic antibodies with desirable features such as: expressibility in a host system, suitability for common protein manufacturing processes while achieving high product purity and yield, and exhibiting high stability during long-term storage conditions. In addition, these therapeutic libraries must also contain antibodies exhibiting in-use characteristics such as low viscosity for injectability at high concentration, long elimination half-lives to reduce dosing frequency, and high bioavailability for conservation of the injected dose. Here, we have described an Antibody-GAN approach to the *in silico* design of monoclonal antibodies which retain typical human repertoire characteristics such as diversity and immunogenicity, while raising the possibility of biasing the libraries *in silico* to achieve other desirable biotherapeutic features.

We experimentally validate our *in silico* approach via phage Fab-display of an initial library of ~100,000 GAN sequences and present the biophysical properties of two example GAN antibodies expressed in CHO. While the biophysical data of the CHO-expressed molecules are not sufficient to indicate any causal effect of the structural differences on the biophysical properties, they show that the molecules are folding appropriately and that they exhibit expected biophysical properties. These results show that the Antibody-GAN is capable of enabling study of large, truly diverse sets of thousands of full-length secreted antibodies, and hundreds of millions of antibody Fabs on phage for biophysical properties. These will provide a real basis to identify causal effect, or lack thereof, of structural properties and sequence on biophysical properties - and that data has the potential to feed *in silico* predictive models that are truly generalized across antibodies.

Ongoing research will be needed to determine precisely which antibody sequence, structure, or biophysical features will bias antibody libraries for developability, quality, and efficacy, as there are many nonlinear pathways for antibody optimization. Existing data sets which interrogate antibody therapeutic developability consist of, in a few cases, hundreds of molecules, but more commonly on the order of tens of antibodies. Deriving from sequence, or even structure, such complex properties as the viscosity or the chemical or thermal stability of an antibody will require far more than hundreds of example molecules. Until now, protein scientists have had to rely on previously-discovered antibodies and their nearby variants, which provide a very small random sampling of the true antibody space. The Antibody-GAN allows us to explore the human antibody space in a rational way. Using transfer learning to bias GANs for given properties, either those calculated on structures from homology modeling, or measured on physically expressed antibodies, we can now begin to understand such questions as how engineering for developability affects key properties like affinity and bio-availability. This provides a mechanism for *in silico* and *in vitro* generation of a much wider range of sequences with intentional biases forming deep, rich training sets for human antibody research.

Recent advances in the protein assay space now provide ultra high-throughput methods in phage or yeast to express and interrogate, for example, the stability of molecules,68 and many more will come. We can now rationally design and create vast experimental antibody data sets for those and future methods, and begin to understand the properties of a developable and effective therapeutic drug.

Our Antibody-GAN approach, as a training-set generation tool, will greatly expand our knowledge of antibody design and behavior. It will also change the way we create therapeutics, by better reproducing properties of *in vivo*-derived antibodies with characteristics that can be tuned to make them better suited as biologics, for production and treatment. Humanoid discovery libraries generated in this way, will provide higher quality treatment and a more rapid and cost-effective response to biological threats and disease targets.

## ACKNOWLEDGMENTS

We would like to thank Lindsay N. Deis for her helpful discussions on library expression and characterization. We would also like to thank Joe Miletich for insightful conversations and Paul J. Sample, Randolph M. Lopez, and Leandra M. Brettner for their constructive comments on the manuscript.

## METHODS

### Training Set Data Sources

Data for the training sets was derived from the Observed Antibody Space (OAS) repository.^58^ Raw nucleotide sequences were automatically translated, classified, and structurally aligned using in-house software (Abacus™). The AHo structure numbering system^59^ was used for structural alignments of the variable regions.

To create the training sets, variable regions were first filtered to remove any sequences which were not classified as human variable regions, by our in-house software Abacus™, and then further cleaned to remove those sequences which contained stop codons, truncations, or insertions. Any sequence which had less than 85% agreement to its closest germline was also removed.

For any paired-chain model, the distinct sequences whose closest germline belonged to the heavy and light germline frameworks of interest were then extracted. These two subsets were randomly sampled and combined during training to create paired sequences.

For any single-chain model, the initial training set contained all represented germlines. If transfer learning was necessary, it was done on an extracted set of sequences whose closest germline belonged to the specific germline of interest.

### Antibody-GAN Development and Training

The Antibody-GAN code was developed in Python. The Keras and Tensorflow deep-learning libraries were primarily used to build and train the Antibody-GAN. The Pandas and Numpy libraries were used to handle any data and training set construction. Other public libraries that were used in the development and analysis of the Antibody-GAN include: Sklearn, Scipy, and Seaborn.

The architecture of the Antibody-GAN is based on the Wasserstein-GAN (WGAN) (with gradient penalty) architecture,^50^ and therefore consists of a generator and a discriminator, which in the WGAN architecture is commonly referred to as a critic. The single-chain network generator (**Figure S1A**) takes as input a noise vector of size 296. This vector is fed into a dense layer, followed by 3 up-sampling and 2D convolutional transpose layers, and a final SoftMax layer to produce a 2D array of size 148×22. This 2D array corresponds to a one-hot-encoded representation of 148 residues and 22 possible amino acids (including deletions and Xs) of a light or heavy chain in an antibody sequence. Antibody sequences aligned by AHo numbering have 149 residues in either chain; to make the network structure simpler, we chose to remove one residue, which appears relatively constant in human repertoire, from each chain during encoding. When decoding, we add this constant residue back in. The discriminator, or critic, takes as input the same 148×22 encoding of an antibody chain and passes it through two 2D convolution layers, followed by a flattening, dense layer and a single-node linear output.

The paired Antibody-GAN architecture (**Figure S2B**) is similar to the single-chain version, except that there are two generators with the same architecture (one per chain). The outputs of each independent-chain generator are concatenated into a 296×22 one-hot-encoded representation of an antibody sequence with both heavy and light chains. It is possible to extend the architecture when training or transfer learning to complex properties that require nonlinear interaction between the two chains. The paired-GAN critic takes as input a 296×22 one-hot-encoded representation of a paired-chain antibody sequence and maintains a similar architecture to the one described above.

**Figure S1C** shows the loss of the generator as well as the discriminator (critic) on fake (generated) and real (training set examples), during training of the single-chain HV3-30 GAN (using a batch size of 128). **Figure S1D** shows the quality assessed by germline framework agreement over training epochs for this model. Training ends when the generated sequences begin showing sufficient quality.

Human monoclonal antibodies have been shown to have higher variability in the heavy chain than the light chain (**Figure 1C** and **Figure S2B**). This may lead to asynchronous optimization of the light chain generator and the heavy chain generator during training in the paired Antibody-GAN, leading to generated heavy chains of higher quality than the light chain. This can be resolved by freezing the layers of the heavy chain generator, once it has reached a state of creating sequences of sufficient quality, and continuing training on the network until the light chain generator has reached a desired quality.

### PFA Set Creation and OAS Set Selection

The IGHV3-30/IGKV3-20 PFA-based sets used in **Figures 1** and **2** were created using the IGHV3-30 and IGKV3-20 training sets extracted from the OAS, which consisted of ~250,000 and ~150,000 sequences respectively. The 100% germline framework for IGHV3-30 was used as a constant framework for all heavy chain PFA sequences, and the 100% IGKV3-20 germline framework was used for all light chains. Each residue in the CDRs (CDR1, CDR2, and CDR3) were then generated using positional frequency analysis; sampling randomly from a distribution representing the frequency of amino acids in the training set, for any given position. 10,000 heavy-chain sequences and 10,000 light chain sequences were created in this manner and then randomly paired together to create a set of 10,000 sequences with full variable regions.

The OAS sets from **Figures 1** and **2** were created by randomly downsampling 10,000 sequences from each of the IGHV3-30 and IGKV3-30 training sets and then pairing together to create a set of 10,000 sequences with full variable regions.

### CDR3 PCA

To perform the PCA analysis, described in **Figure 1**, the aligned CDR3 of a given antibody was one-hot encoded into a vector representation. A 2-component PCA model was built, using the sklearn library, on those one-hot encoded vectors from all sequences of the OAS set, the PFA set, and the base GAN set (totaling 30,000 samples). Heavy chain and light chain models were built and trained separately.

### Antibody-GAN Biasing Sources

#### CDR H3

Our in-house software, Abacus™, was used to assess the length of the CDR H3 from any training set, GAN-generated set, or PFA-generated set.

#### Calculated Immunogenicity

MHCII is a polymorphic transmembrane protein that binds and presents fragments of foreign, extracellular proteins to T-cell receptors (TCRs) to initiate an adaptive immune response. The MHCII binding score reported in **Figure 2B** is a composite metric intended to quantify the immunogenicity risk in a sequence based on whether its constituent peptides are predicted to bind strongly and promiscuously to MHCII proteins. The quality of this metric depends on the selection of an accurate peptide-MHCII binding predictor and a reasonable method for aggregating predictions across the peptide fragments in a sequence and across allelic variants of MHCII.

We developed a machine learning algorithm for peptide-MHCII binding affinity prediction, trained on the peptide-MHCII binding affinity data set used to train NetMHCIIpan-3.2 and reported by Jensen et al.^54^ Several machine learning algorithms have been developed that outperform traditional matrix-based approaches to peptide-MHCII binding affinity prediction, including NetMHCII-pan and, more recently, MARIA.^40,54^ We use our in-house MHCII binding predictor for ease of integration with our other sequence analysis tools and based on favorable accuracy comparisons with published benchmarks (not shown in the present report). **Figure S5A** shows that predictions from our models are generally correlated with the “IEDB recommended” algorithm for peptide-MHCII binding prediction.^60^

To calculate a sequence MHCII binding score, we first break the sequence into each of its constituent 15mer peptide fragments (sliding window of 15, stride of 1). For each 15mer, we use allele-specific models to predict the binding affinity to 8 common allelic variants of MHCII (those encoded by HLA alleles DRB1*0101, DRB1*0301, DRB1*0401, DRB1*0701, DRB1*0801, DRB1*1101, DRB1*1301, and DRB1*1501). This set of alleles was also used in the pioneering MHCII binding risk-reduction work of de Groot and Martin.^7^ We convert the binding affinities into z-scores for each allele using the mean and standard deviation of affinities predicted for a large reference set of 15mers randomly selected from the human protein sequences stored in UniProt.^69^

We take the median z-score across alleles for each 15mer, and sum the positive median z-scores across the sequence to get the final MHCII binding score. The median is an appropriate aggregation because a peptide fragment that binds several MHCII variants poses an immunogenic risk to a larger population of patients than a fragment that binds to only one. Dhanda et al., creators of the protein deimmunization engine on the IEDB website, also aggregate MHCII binding scores across alleles using the median.^53^ We ignore negative scores in our sum across the sequence because peptides that certainly don’t bind to MHCII (large negative scores) should not offset peptides that bind MHCII tightly (large positive score). **Figure S5B** shows the fraction of putative MHC binding peptides for all unique 15mers in each sequence set described in **Figure 2B**. The low-immunogenicity set (GAN −I) has fewer MHCII-binding peptides than other sets, suggesting the GAN learns which 15mers to avoid regardless of our sequence-score abstraction.

#### Structure Modeling, Calculated Isoelectric Point (pI), and Negative Patch Surface Area

Structure models were calculated as Fab structures using the antibody modeling tool within the Molecular Operating Environment (MOE, Chemical Computing Group, Montreal, Canada). Fab structures were used rather than Fvs in order to generate more accurate Fv surface patches in the presence of constant domains. The pIs were calculated using the Ensemble Isoelectric Point method in the Protein Properties tool within MOE called as an SVL method. The electronegative patch sizes were calculated using the Protein Patches method as an SVL call within MOE with the Hydrophobic Min Area (p_hminarea) changed from the default setting of 50 to 30 Å^2^ and the Charge Cutoff (p_qcutoff) changed from the default setting of 40 to 20 Å^2^.

### GAN-Library Sequence Selection

Human repertoire contains a small subset of sequences which have missing residues, non-standard cystines, non-standard N-linked glycosylation sites, or potential N-linked glycosylation sites. Sequences with these properties were not pulled from the training set and are therefore also represented by a small subset in the GAN libraries. For our phage library, we filtered out any sequences generated by the GAN which had any of these properties, before selecting final sequences.

### Phage Expression of GAN-Library

#### Bacterial strains

*Escherichia coli* One Shot™ TOP10 cells (F- *mcrA* Δ(*mrr-hsd*RMS-*mcr*BC) Φ80*lac*ZΔM15 Δ *lac*X74 *rec*A1 *ara*D139 Δ(*araleu*)7697 *gal*U *gal*K *rps*L (StrR) *end*A1 *nup*G) were purchased from Thermo Fisher Scientific and used for phagemid DNA cloning. E. cloni® 10G electrocompetent cells (F− *mcrA* Δ(*mrr-hsd*RMS-*mcr*BC) *end*A1 *rec*A1 Φ80*dlac*ZΔM15 Δ*lac*X74 *ara*D139 Δ(*ara,leu*)7697*gal*U *gal*K *rps*L *nup*G λ- *ton*A (StrR)) were purchased from Lucigen Corporation and were also used for phagemid DNA cloning. *E. coli* SS320 electrocompetent cells (*F’[proAB lacI*^*q*^*Z ΔM15 Tn10 (Tet*^*R*^*)] araD139 Δ(ara-leu)7696 galE15 galK16 Δ(lac)X74 rpsL (Str*^*R*^*) hsdR2 (r*_*K*_*– m*_*K*_*+) mcrA mcrB1*) were purchased from Lucigen Corporation and used as the host for phage library production.

#### Cloning

The phagemid pADL-20c (Antibody Design Labs) was used for construction of the GAN sub-libraries and was modified for expression of Fab antibody fragments as N-terminal pIII fusion proteins in *E. coli*. This vector utilizes the bacterial pectate lysate (pelB) signal sequences for periplasmic translocation of fusion proteins, along with an ampicillin resistance gene for growth and selection in transformed *E. coli*. A hexahistidine tag and a FLAG tag was added to the C-terminus of the CH1 and kappa constant domains, respectively, and the amber stop codon upstream of gIII was removed to allow expression of the fusion protein in SS320 host cells.

Synthetic gene fragments encoding variable heavy and light chains were first amplified individually using PCR primers containing 22 base pairs of sequence complementary to the phagemid backbone. Next, PCRs were pooled by germline and assembled sequentially into the phagemid using NEBuilder® HiFi DNA Assembly Master Mix (New England Biolabs). Transformations were performed using One Shot™ TOP10 or E. cloni 10G® cells, and the resulting phagemid DNA was purified using ZymoPURE™ II Plasmid Midiprep Kit (Zymo Research).

#### Phage library production

*E. coli* SS320 host cells were electroporated as described by the manufacturer using 250 ng of each sub-library DNA. An aliquot of each transformation was plated on 2xYT agar plates supplemented with 100 μg/mL carbenicillin and 2% glucose and incubated overnight at 30°C. The resulting colonies were used for estimating library size and for sequencing the variable heavy and light chains using colony PCR. The remainder of the transformation was used to inoculate 2xYT-CG (2xYT broth containing 50 μg/mL carbenicillin and 2% glucose) at an OD_600 nm_ of 0.07 and incubated with shaking at 250 rpm and 37°C until an OD_600 nm_ ~ 0.5. The cultures were then infected with M13KO7 helper phage (Antibody Design Labs) at a multiplicity of infection (MOI) of 25 and incubated at 37°C without shaking for 30 minutes, followed by shaking at 200 rpm for 30 minutes. Cultures were centrifuged followed by medium replacement in 2xYT-CK (2xYT supplemented with 50 μg/mL carbenicillin and 25 μg/mL kanamycin). After overnight incubation at 30°C and 200 rpm, the phage particles were purified and concentrated by PEG/NaCl precipitation and resuspended in PBS containing 0.5% BSA and 0.05% Tween-20. Phage concentration was determined using a spectrophotometer, assuming 1 unit at OD_268 nm_ is equivalent to 5×10^12^ phage/mL. PEG-precipitated phage from each GAN sub-library was normalized to 1×10^13^ phage/mL and serially diluted 10-fold in 2% non-fat dry milk in PBS, in duplicate, for use in the polyclonal phage ELISA.

#### Monoclonal phage production

Single clones harboring a functional Fab fusion protein were inoculated into 500 μL 2xYT-CTG (2xYT broth supplemented with 50 μg/mL carbenicillin, 15 μg/mL tetracycline, and 2% glucose) and cultivated in 96 deep-well plates overnight at 37°C with rigorous shaking. 5 μL of the overnight cultures was then transferred to new deep-well plates containing 100 μL 2xYT-CTG and incubated at 37°C with rigorous shaking until an OD_600 nm_ ~ 0.5. M13KO7 helper phage was added to each well at MOI25, and plates were incubated without agitation at 37°C for 1 hour before medium replacement to 2xYT-CK and overnight incubation with rigorous shaking at 30°C. Phage supernatants were harvested after centrifugation and diluted 1:1 in 2% non-fat dry milk in PBS for use in the monoclonal phage ELISA.

#### Phage ELISA

The amount of Fab displayed on phage was determined using ELISA. Briefly, 96-well MaxiSorp® assay plates (Nunc) were coated overnight at 4°C with anti-human Fab (Millipore Sigma) diluted 1:500 in PBS, then blocked in PBS containing 1% BSA for 1 hour at room temperature (RT). Diluted phage preparations were added and allowed to incubate for 1 hour at RT before captured virions were detected using a 1:5000 dilution of anti-M13-HRP (Santa Cruz Biotechnology) for 1 hour at RT. All interval plate washes were performed 3 times in PBST (PBS supplemented with 0.1% v/v Tween-20). ELISAs were developed by addition of TMB solution (Thermo Fisher Scientific) and quenched using 10% phosphoric acid. Absorbance was read at A_450 nm_. Phage supernatant from non-transformed *E. coli* SS320 host cells was included as a negative control.

### Selected CHO-Expressed Molecules

CHOK1 Glutamine Synthetase (GS) knockout host cells (Horizon Discovery, Cambridge, United Kingdom) were maintained in CD OptiCHO (Thermo Fisher Scientific, Waltham, MA) containing 4 mM glutamine. Cells were cultured as previously described.^70^

The light chains (LC) and heavy chains (HC) containing appropriate signal peptides were cloned into an in-house proprietary bicistronic PiggyBac transposon expression vector^70^ in a sequential, 2-step manner using Gibson assembly. Successful insertion of the intended coding sequences was confirmed by Sanger DNA sequencing. Plasmid DNA was purified using a conventional silica-based low endotoxin Zymo Research kit (Irvine, CA).

Cells, DNA and RNA were added to BTX 25 – multi-well electroporation plates (Harvard Bioscience, Holliston, MA) using a Tecan Freedom EVO (Mannedorf, Switzerland) liquid handler. For each transfection, 2.4E6 cells were spun down and resuspended in 150uL of PFCHO medium (Sigma-Aldrich, St. Louis, MO). 7.5ug of DNA and 2.5ug of pJV95 transposase RNA were added to the cells, then electroporated at 3175 uF capacitance, 290 V voltage, 950 Ω resistance in an ECM 630 electro manipulator coupled to a HT 100 high throughput adaptor (BTX, Holliston, MA). Both molecules were transfected in triplicate. Cells were transferred to 2mLs of non-selective medium in a 24 deep well plate (DWP) shaking at 220rpm at standard growth conditions and cultured for 2 days prior to selection. After two days, cells were counted on a guava flow cytometer (Luminex, Austin, TX); plates were spun down and resuspended in 2mLs of selective CD OptiCHO medium. Cells were counted and passaged every four to five days thereafter.

Thirteen days after selection started and viability was >90%, cells were seeded into proprietary production medium at 8 × 105 c/mL in 3mLs in 24 DWPs under standard growth conditions. On days 3, 6 and 8, cells were fed with 5% of the starting volume with Cell Boost7a and 0.5% Cell Boost7b (Hyclone GE Healthcare Life Sciences). Cell counts and glucose was measured as previously described,^70^ On day 8, 50% glucose was supplemented to a final concentration of approximately 10g/L. On day 10, cells were counted, spun down and filtered by centrifugation onto 24-deep well filter plates (Thomson, Oceanside, CA). Titer was sampled by Ultra High-Performance Liquid Chromatography (UHPLC) Protein A affinity. Replicate wells were pooled together for Protein A purification.

### Biophysical Validation of CHO-Expressed Molecules

#### Sample preparation

Samples were buffer exchanged against 10 diavolumes of 20mM sodium chloride, 150mM sodium chloride, pH 7.1 (PBS) using a centrifugal filter with a 30kDa molecular weight cut off (Amicon). After buffer exchange, samples were normalized to 1 mg/mL using a Lunatic protein concentration plate format instrument (Unchained Labs).

#### Differential Scanning Fluorimetry

Thermal transition temperature(s) and weighted shoulder scores were determined by DSF according to the method previously described (Kerwin, 2019).

#### Self Interaction Nanoparticle Spectroscopy

SINS measurements were performed according to the method previously described (Liu, 2013) Briefly, gold nanoparticles (Ted Pella) were conjugated overnight with an 80:20 ratio of anti-human and anti-goat antibodies (Jackson Immuno Research). Unreacted sites were blocked using an aqueous 0.1% (w/v) polysorbate 20 solution. Conjugated gold nanoparticles were then concentrated by centrifugation and removal of 95% of the supernatant. Analysis was carried out in PBS (20 mM phosphate, 150 mM NaCl, pH 7.1) at a protein concentration of 0.05 mg/mL reacted with 5 ul of concentrated conjugated gold nanoparticles. After a 2 hour incubation, absorbance spectrum from 400-600 nm was collected using a Spectrostar Nano plate reader at 2nm steps. The wavelength maximum of the spectrum peak is reported.

#### Relative Solubility

Solubility was assessed according to the method previously described (Kerwin, 2019). Analysis was done in PBS buffer (20 mM sodium phosphate and 150 mM sodium chloride pH 7.1) and a final PEG 10,000 concentration ranging from 0% to 12%. Remaining soluble protein after PEG incubation is reported.

#### Size Exclusion High Performance Liquid Chromatography

Size exclusion high performance liquid chromatography (SEC) was performed on a Dionex UltiMate 3000 HPLC System using a Waters XBridge Protein BEH SEC 200Å, 3.5 μm column and a diode array detector collecting at 280 nm. Separation was achieved under native conditions with a 100 mM sodium phosphate, 250 mM sodium chloride, 10% acetonitrile v/v mobile-phase buffer at pH 6.8.

## SUPPLEMENTARY

**Figure S1.** Single-chain and paired-chain Antibody-GAN architectures and training. **A)** Detailed architecture of single-chain Antibody-GAN; **B)** Detailed architecture of full paired-chain Antibody-GAN; **C)** Single-chain HV3-30 Generator and Discriminator (Critic) model losses over training epochs; **D)** Quality of generated HV3-30 sequences over training epochs assessed by germline agreement to framework.

**Figure S2.**
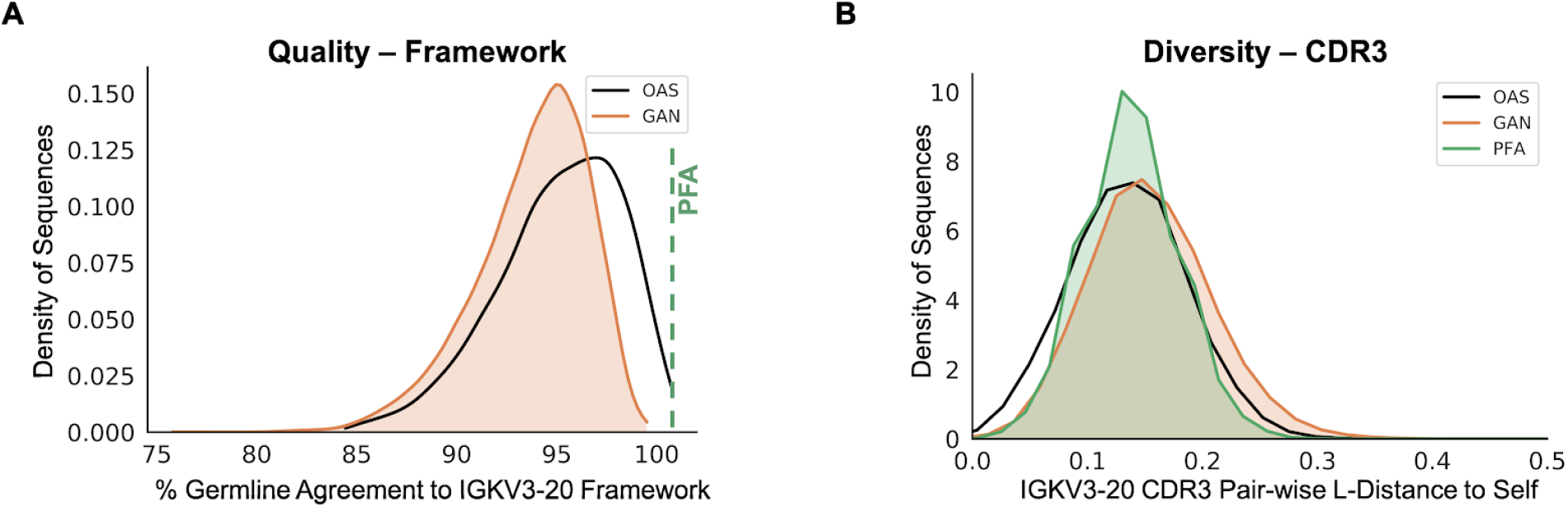
Kappa light chain (KV3-20) quality and diversity. **A)** Representative percent germline agreement for KV3-20 framework residues from the training (OAS) and the base, unbiased GAN (GAN) for 10,000 generated sequences per set **B)** Representative diversity in light-chain KV3-20 CDR3, as Levenshtein distance calculated within sets of 10,000 sequences from each of: the OAS training set, GAN-generated molecules, and CDR PFA-generated molecules.

**Figure S3.**
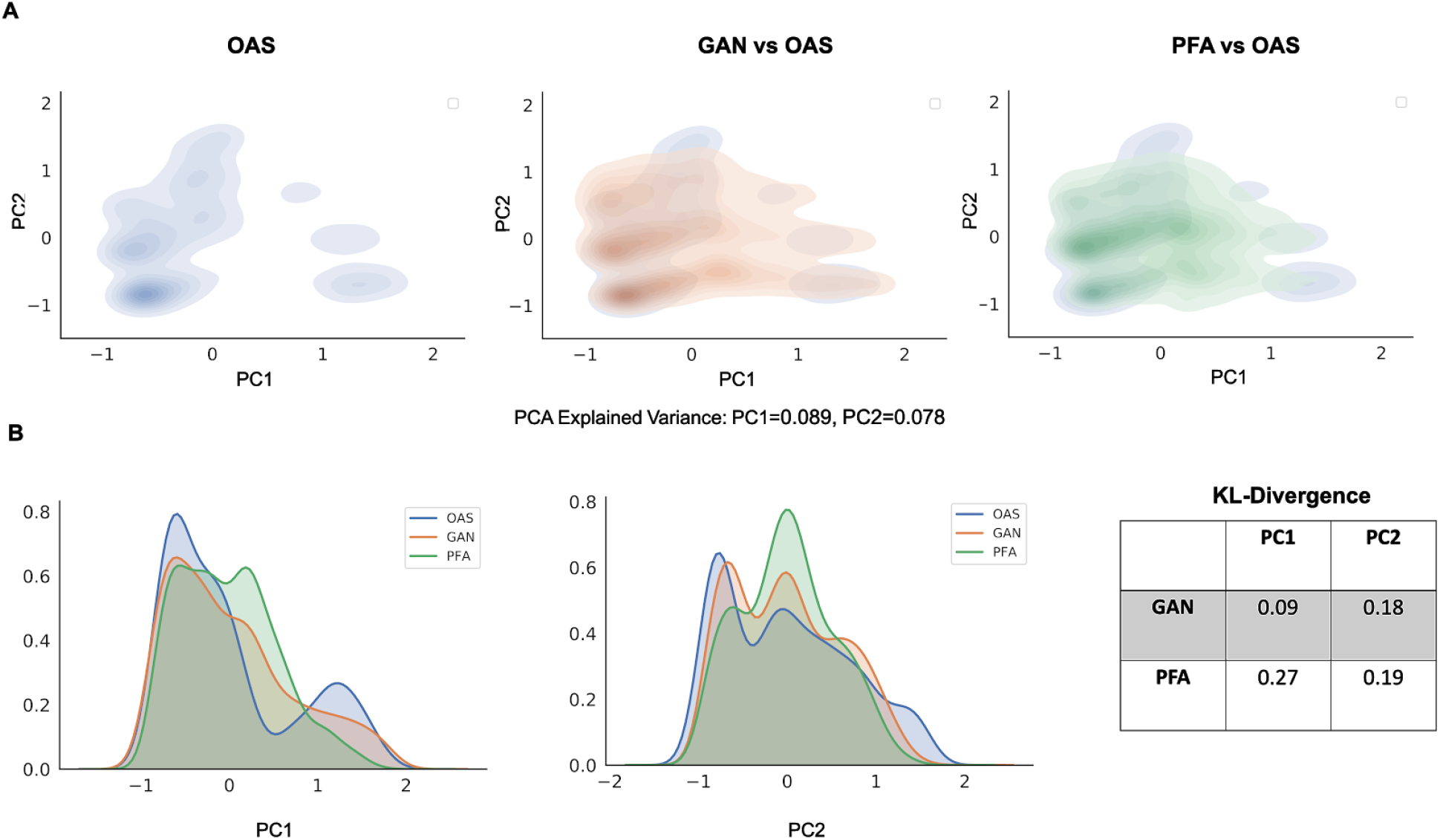
Raw CDR3 PCA score comparisons for KV3-20. **A)** Quality and diversity of light (kappa) chain CDR3 as captured in the scores of a principal component analysis (PCA) with explained variance of 9% for PC1 and 7% for PC2. The GAN set shows more similar coverage in the PCA space to human repertoire (OAS) than PFA. Both PFA and GAN capture similar densities compared to OAS; **B)** The KL-divergence of each principal component for GAN and PFA, compared to OAS, shows lower divergence of GAN from OAS in PC1 and a similar low divergence for both in PC2.

**Figure S4.**
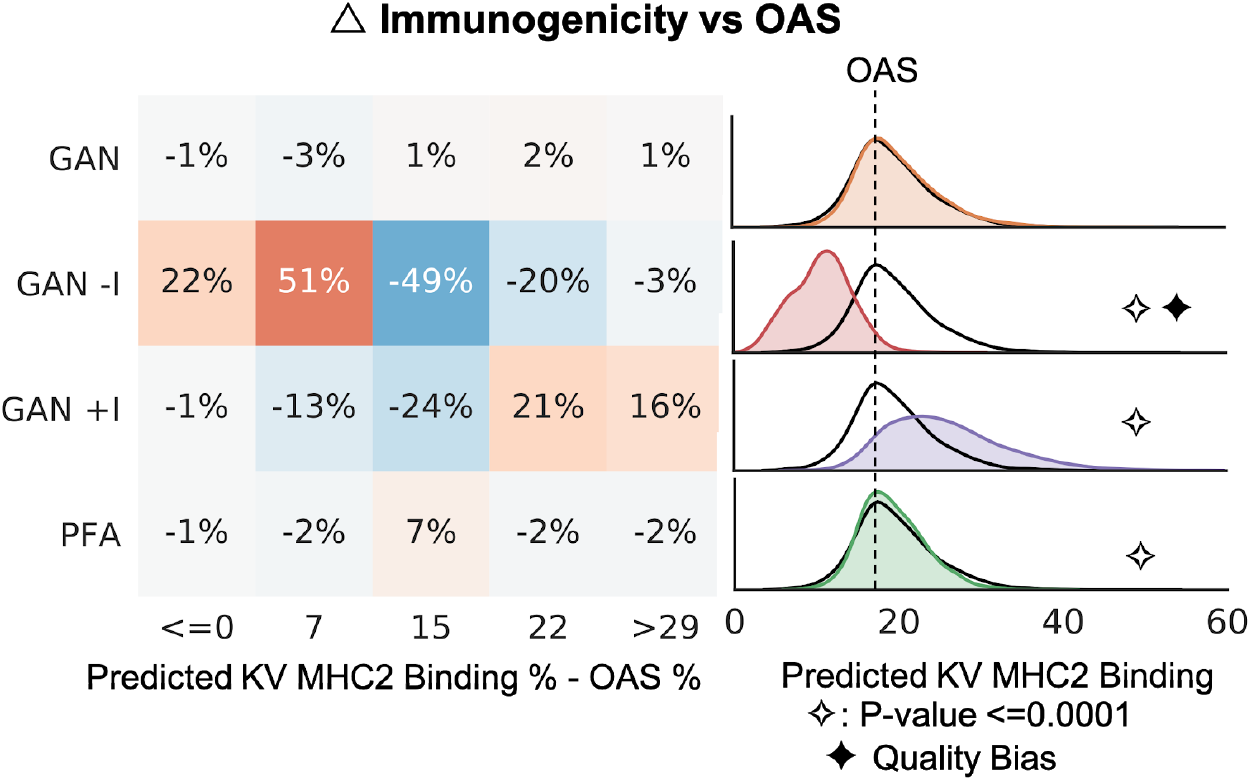
Light chain (KV3-20) MHCII biasing. Representative predicted light chain (KV3-20) MHCII binding, as compared to OAS, for the unbiased Antibody-GAN (GAN), Antibody-GANs transfer learned to lower (GAN −I) and to higher (GAN +I) predicted MHCII binding, and PFA-generated (PFA) molecules; showing the GAN set to be statistically indistinguishable from OAS (human repertoire), while PFA diverges slightly but significantly. The GAN −I was intentionally biased by approximately a 73% shift to lower predicted MHCII binding, while the GAN +I set managed an intentional bias of ~37% to higher predicted MHCII binding.

**Figure S5.**
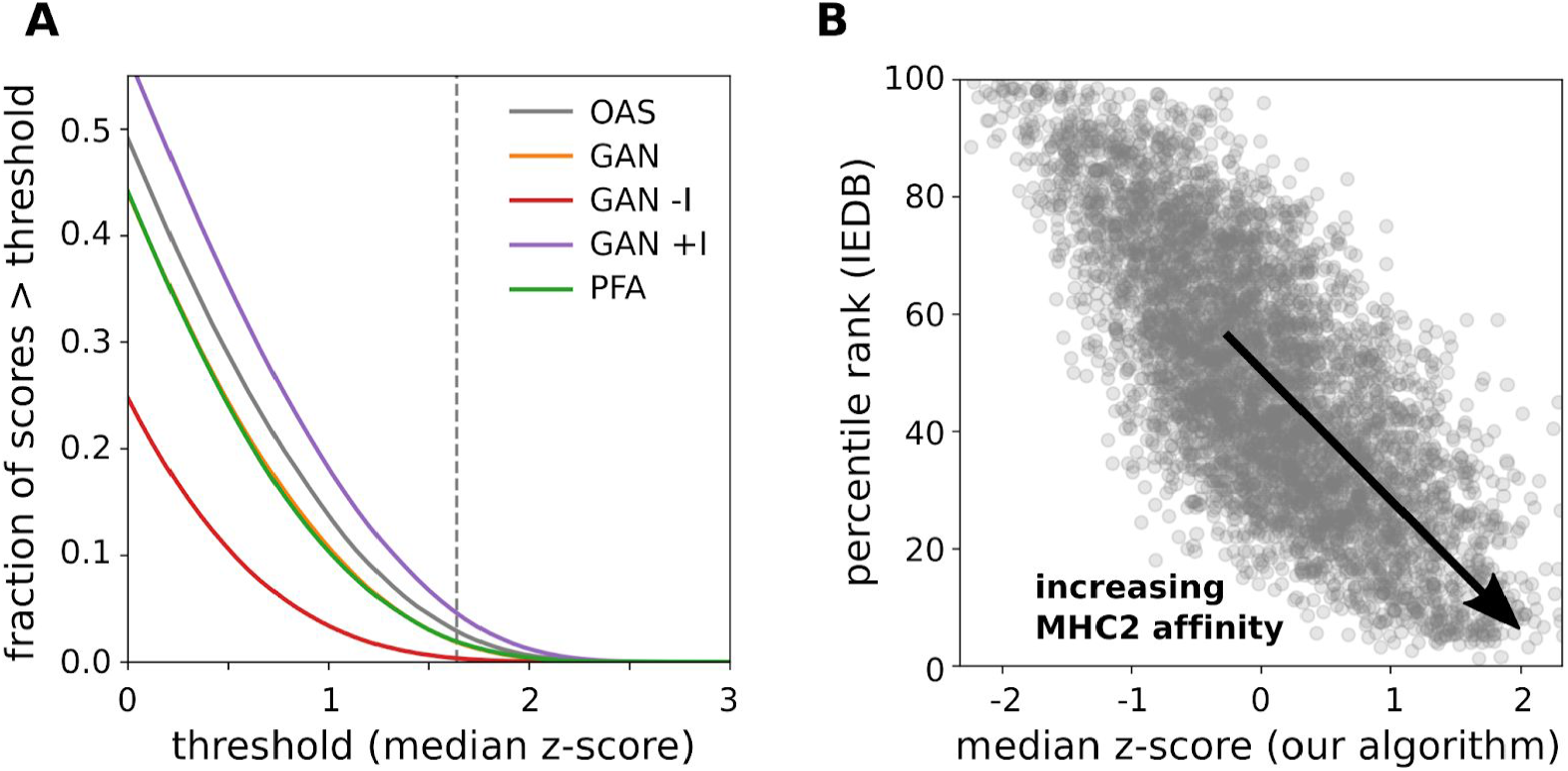
MHC-II binding score bias is robust to changes in metric or predictor. **A)** The fraction of unique 15mers classified as promiscuous MHC-II binders using median z-score thresholds between 0 and 3, for the OAS (gray), GAN (orange), GAN −I (red), GAN +I (purple), and PFA (green) data sets compared in Figure 2b. Less than 1% of the unique 15mers in the GAN −I data set exceed a z-score of 1.64 (dashed line), corresponding to the top 5th percentile, and used in other works as a threshold for identifying potential “immunogenicity hotspots”. **B)** Percentile rank assigned by the “IEDB recommended 2.22” prediction method vs. z-score assigned by our in-house MHC-II binding algorithm, calculated as the median across the same 8 allelic variants of MHC-II described in the main text for a random sample of 5000 peptides from our OAS reference set. Our method is generally correlated with the IEDB recommended MHC-II binding algorithm, but we choose our own method for its ease high-throughput use and favorable accuracy on benchmark data sets (comparison not shown).

**Figure S6.**
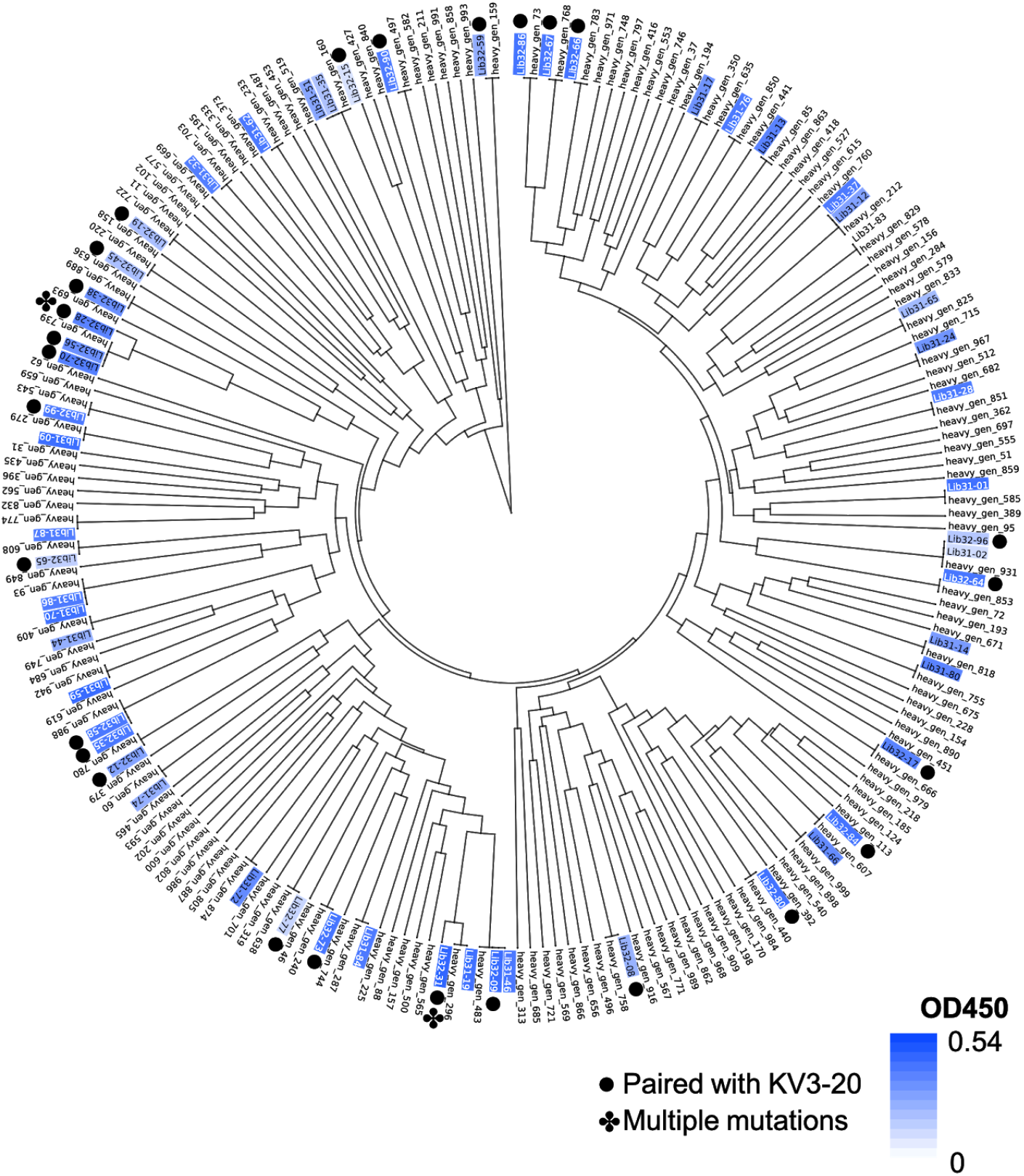
Clading of selected monoclonal phage sequences on HV1-2. Clading of variable region protein sequences of select phage colonies expressing the HV1-2 library as determined by DNA sequencing. Blue highlighting indicates degree of expression level of selected Fabs as determined by ELISA. Sequences that were paired with KV3-20 light chains(●) are as marked, otherwise they were paired with KV1-39 light chains. Sequences with multiple amino acid substitutions(✤) as compared to their intended GAN design are as marked.

**Figure S7.**
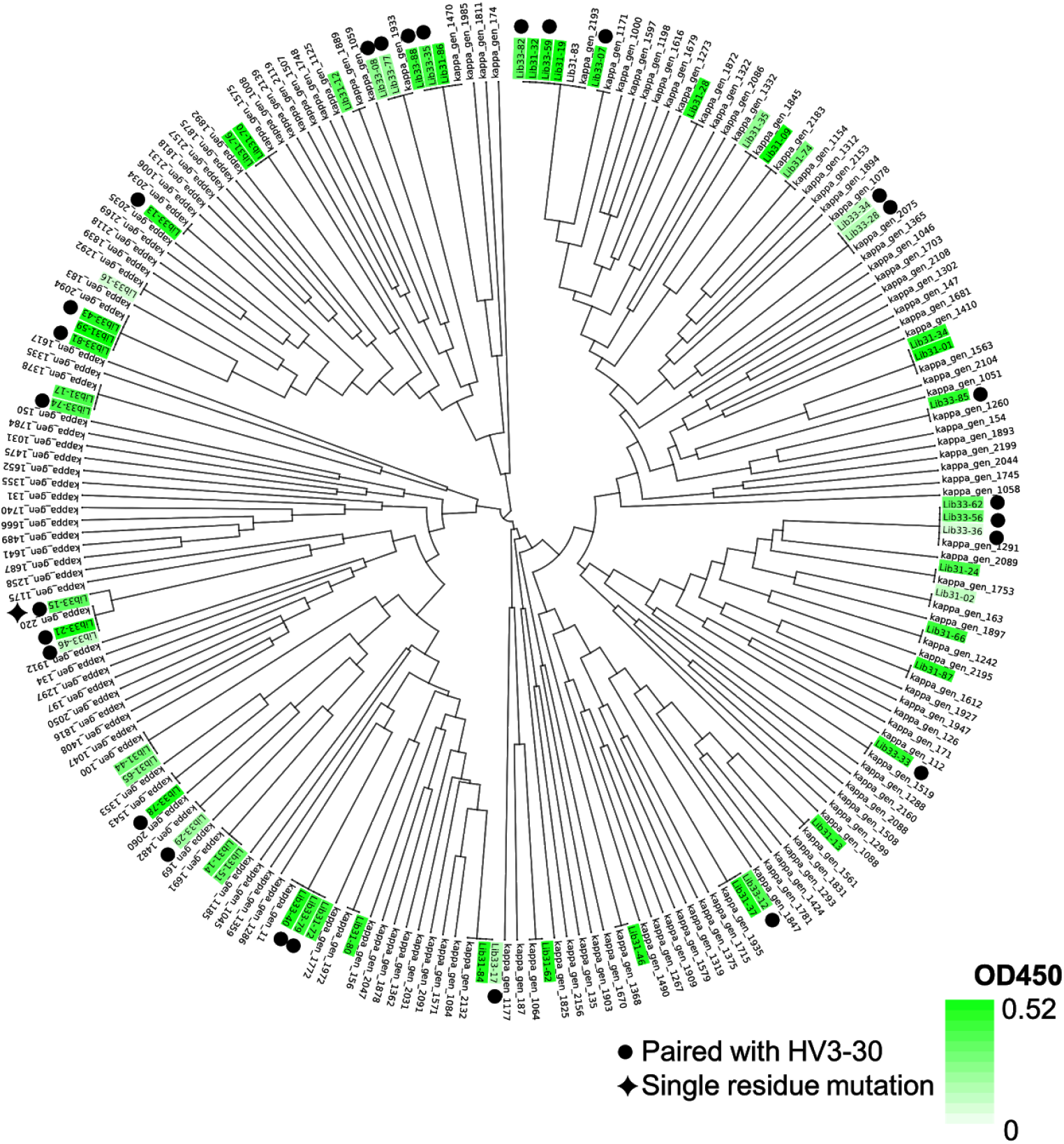
Clading of selected monoclonal phage sequences on KV1-39. Clading of variable region protein sequences of select phage colonies expressing the KV1-39 library as determined by DNA sequencing. Green highlighting indicates degree of expression level of selected Fabs as determined by ELISA. Sequences that were paired with HV3-30 heavy chains(●) are as marked, otherwise they were paired with HV1-2 heavy chains. Sequences with single amino acid substitutions(✦) as compared to their intended GAN design are as marked.

**Figure S8.**
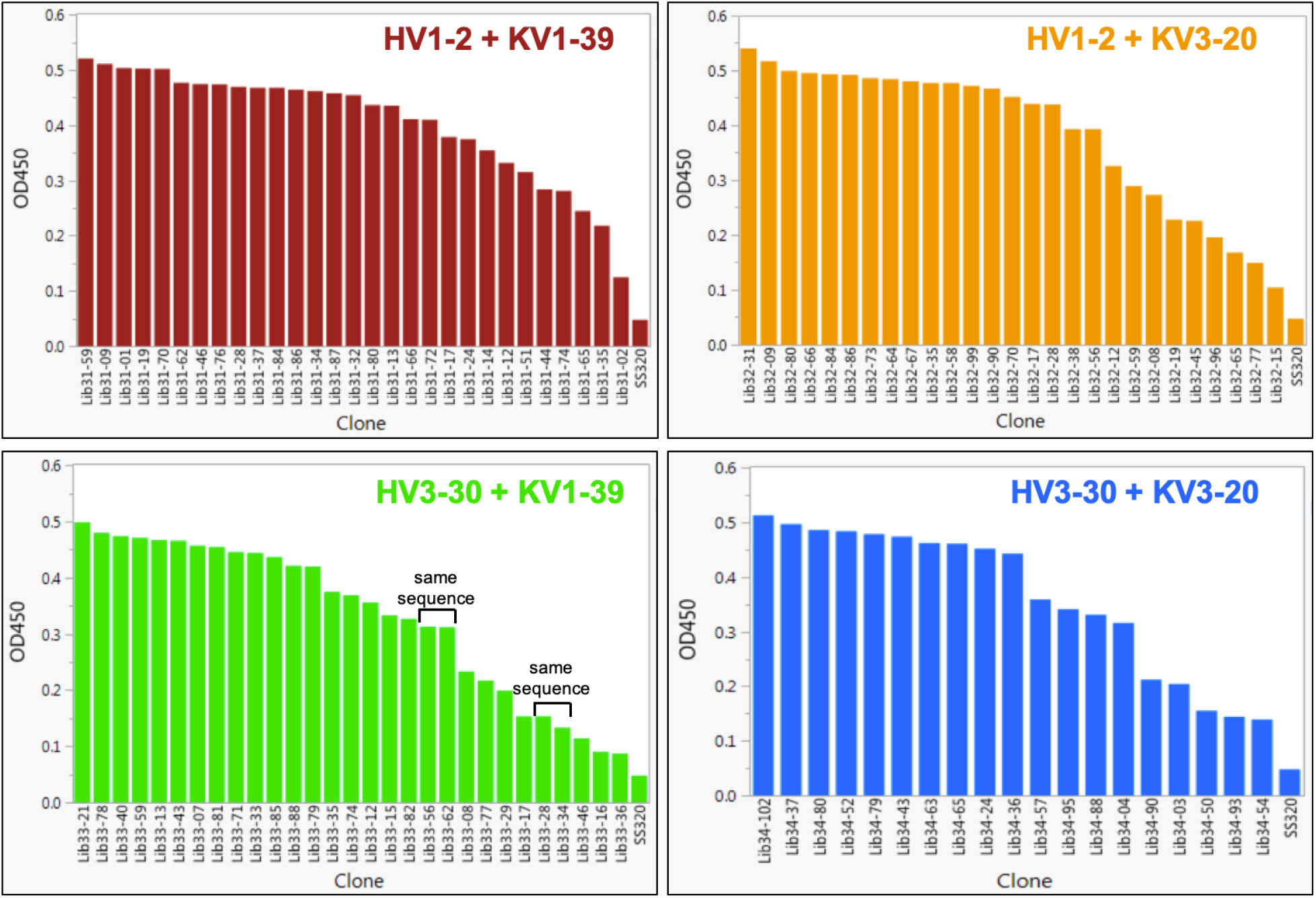
Monoclonal phage expression results with controls. Expression level of selected Fabs as determined by ELISA from each of the 4 GAN sub-libraries, showing selected Fabs having higher OD450 than control phage 2230 not expressing Fab. Two pairs of colonies were found to have identical heavy and light chains and showed very similar ELISA results.

**Figure S9.**
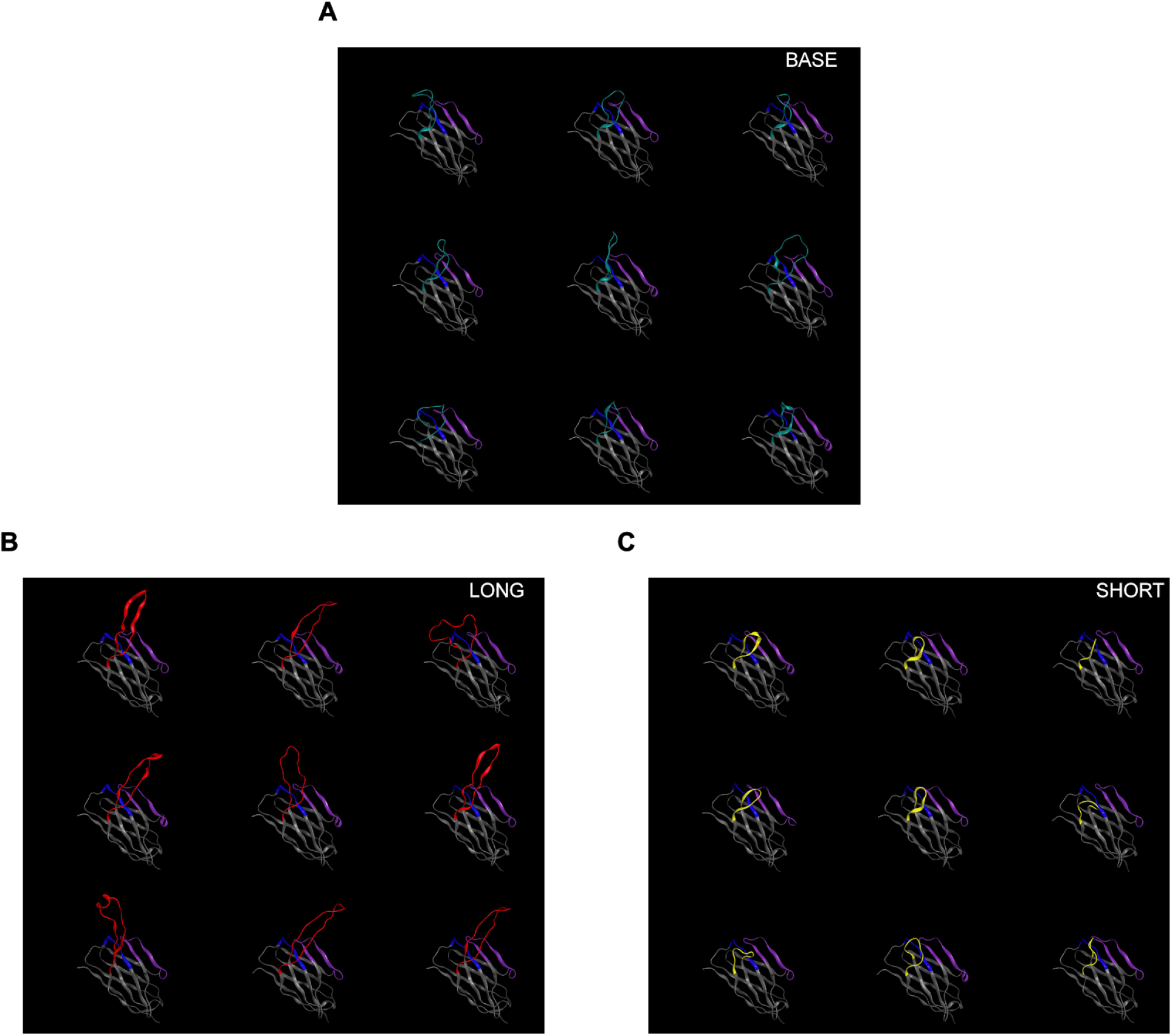
Snapshot of molecules from base GAN and CDR3-biased GANs during training. Snapshots of 9 generated molecules during training of **(A)** the base GAN, **(B)** the base GAN transfer-learned to long CDR3s, and **(C)** the base GAN transfer-learned to short CDR3s. Loops colored in teal indicate a CDR3 of length >= 11 amino acids long and <= 20 amino acids long. Loops colored in red indicate CDR3s of length >20, and those colored in yellow denote a CDR3 of length <11 amino acids.

## Notes

### Competing Interest Statement

All authors are employed by Just - Evotec Biologics, Inc whose business includes biotherapeutic discovery and development.

### Summary of Updates

Supplementary Figure S9 added; 2 movies associated with Figure S9 showing antibody structures during training and transfer learning of Antibody-GAN on 9 example seeds.

